# Adaptive optical correction for *in vivo* two-photon fluorescence microscopy with neural fields

**DOI:** 10.1101/2024.10.20.619284

**Authors:** Iksung Kang, Hyeonggeon Kim, Ryan Natan, Qinrong Zhang, Stella X. Yu, Na Ji

**Affiliations:** Department of Neuroscience, University of California, Berkeley, CA 94720, USA; Applied Science and Technology Graduate Program, University of California, Berkeley, CA 94720, USA; Department of Electrical Engineering and Computer Science, University of Michigan, Ann Arbor, MI 48109, USA; Department of Electrical Engineering and Computer Science, University of California, Berkeley, CA 94720, USA; Department of Physics, University of California, Berkeley, CA 94720, USA; Helen Wills Neuroscience Institute, University of California, Berkeley, CA 94720, USA; Molecular Biophysics and Integrated Bioimaging Division, Lawrence Berkeley National Laboratory, Berkeley, CA 94720, USA

## Abstract

Adaptive optics (AO) restore ideal imaging performance in complex samples by measuring and correcting optical aberrations, but often require custom-built microscopes with carefully aligned wavefront sensing/shaping devices and can be susceptible to sample motion. Here we describe NeAT, a computational framework using neural fields for AO two-photon fluorescence microscopy. NeAT estimates wavefront aberration and recovers sample structure from a 3D image stack without requiring external datasets for training. Incorporating motion correction in learning and correcting conjugation errors commonly found in commercial microscopes, NeAT is designed for deployment in biological laboratories for *in vivo* imaging. We validate NeAT’s performance using a custom-built microscope with a wavefront sensor under varying signal-to-noise ratios, aberration, and motion conditions. With a commercial microscope, we demonstrate real-time aberration correction for *in vivo* morphological and functional imaging in the living mouse brain, with NeAT improving signal and accuracy of glutamate and calcium imaging of synapses and neurons.

## Introduction

Fluorescence imaging of living biological organisms provides mechanistic insights into their physiology. Two-photon (2P) fluorescence microscopy is an essential tool for live imaging, probing structure and function at subcellular resolution deep within complex tissues^1^. However, as 2P excitation light propagates through tissue, its wavefront accumulates optical aberrations from refractive index mismatches, leading to reduced fluorescence signal, resolution, and contrast. When these sample-induced aberrations are measured and corrected, the excitation light can form a diffraction-limited focus, increasing fluorescence signal and improving the accuracy of structural and functional characterization.

Adaptive optics (AO)^2–6^ measure aberration and correct it with wavefront-shaping devices, such as deformable mirrors (DM) and liquid-crystal spatial light modulators (SLM). AO methods can be grouped into direct wavefront sensing methods, which use a wavefront sensor for aberration measurement, and indirect wavefront sensing methods, which include approaches utilizing machine learning for wavefront estimation^7–12^.

Regardless of aberration measurement scheme, AO methods are generally developed for and deployed on custom-built microscopes, in which individual optical components are carefully conjugated and aligned to ensure optimal imaging and correction performance. However, microscopes in a general laboratory setting often have imperfect conjugation and misalignment of optical components, with commercial microscopes additionally suffering from limited access and adjustability of their optical paths. Furthermore, sample motion during *in vivo* imaging leads to artifacts that degrade aberration measurement accuracy, a problem that can be particularly severe for deep tissue imaging as well as for indirect wavefront sensing methods that utilize serial measurement of images and signals^4^.

Here, we describe NeAT, **<u>Ne</u>**ural fields for **<u>A</u>**daptive optical **<u>T</u>**wo-photon fluorescence microscopy. It utilizes neural fields to represent a sample’s three-dimensional (3D) structure and incorporates computational architectures to enhance AO performance for imperfect microscopes and living samples. By incorporating an image-formation model for two-photon fluorescence microscopy that accounts for both aberration and sample motion as a physics prior, NeAT accurately estimates aberration from a single fluorescence image stack without requiring external datasets for training, even in the presence of motion artifacts. NeAT also corrects for conjugation errors in the microscope system, ensuring that the corrective phase pattern displayed on a wavefront-shaping device accurately cancels out aberration after propagation through imperfectly conjugated and misaligned optics. Lastly, NeAT jointly recovers the sample’s 3D structure with aberration. In scenarios where additional imaging with aberration correction is not needed, NeAT eliminates the need for corrective devices, further reducing system cost and complexity.

The paper is structured as follows. First, we implement NeAT in a perfectly conjugated 2P fluorescence microscope equipped with a wavefront sensor for direct wavefront sensing (DWS). We then compare the performance of NeAT with ground-truth aberration measurement by DWS both *in vitro* and *in vivo*, and determine its performance limits in terms of signal-to-noise ratio (SNR), aberration severity, and sample motion. Next, we implement NeAT in a commercial microscope with imperfect conjugation and evaluate its real-time aberration correction performance for *in vivo* morphological and functional imaging within the living mouse brain.

## Results

### NeAT, a general-purpose AO framework in 2P fluorescence microscopy using neural fields

NeAT is designed to jointly estimate wavefront aberration and recover sample structure from an input 3D 2P fluorescence image stack (**Fig. 1**). It utilizes neural fields – implicit functions represented by a coordinate-based neural network across spatial coordinates^14^ – to represent sample structure (**Fig. S1a**). NeAT also incorporates a mathematical image-formation model for 2P fluorescence microscopy into the learning process, which involves aberration and structural estimation, as well as motion correction through learnable image transformations. During the learning process, NeAT aims to reproduce an image stack closely resembling the input by iteratively adjusting its parameters. This process requires no external supervision.

**Figure 1.**
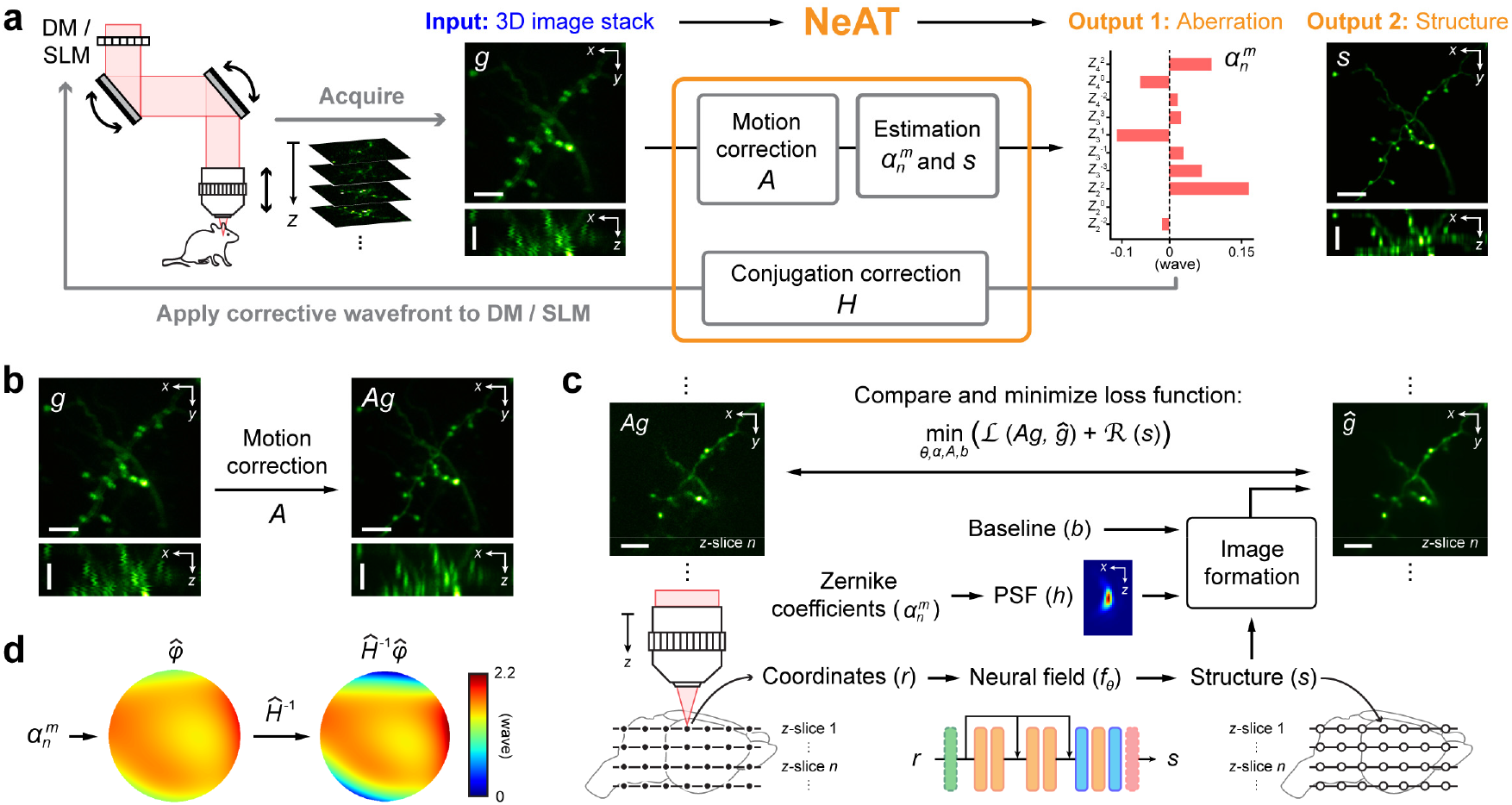
NeAT estimates aberration and recovers structure from a 3D input image stack. **(a)** Schematic of NeAT’s function and integration into an adaptive optics (AO) imaging pipeline. Lateral (*xy*) and axial (*xz*) maximum intensity projections (MIPs) of a 3D two-photon (2P) fluorescence image stack (*g*) as the input to NeAT. The learning process of NeAT corrects for sample motion, and outputs an estimated aberration (Zernike coefficients 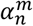) and sample structure (*s*). Aberration output is then used to calculate a corrective wavefront to be applied to a deformable mirror (DM) or liquid-crystal spatial light modulator (SLM) for aberration correction. If present, microscope conjugation errors are measured by NeAT and compensated for before applying corrective wavefront. Scale bar: 5 µm. **(b)** If present, sample motion artifacts in *g* are corrected by applying learnable transformations *A*. **(c)** NeAT optimizes network weights (*θ*), *ĝ* · *ĝ*, and *A* to minimize loss function ℒ (*Ag, ĝ*) + ℛ (*s*) that compares the input image stack *Ag* with an image stack 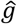is computed from a structure represented by a neural field (*s* = *f*_θ_ (***r***)) and a 3D point spread function calculated from 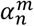 (PSF *h*; axial MIP shown). **(d)** If present, conjugation errors are estimated as *Ĥ* and compensated for by applying *Ĥ* ^−1^ to the corrective phase pattern. 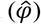 Scale bars: 5 µm.

The input for NeAT is an image stack (*g*) acquired through *z*-axis scanning (**Fig. 1a**). Artifacts caused by sample motion (e.g., body movement, breathing, and heartbeat) in the *z* stack, if present, are corrected by a set of affine transformations (*A*) whose parameters are optimized during the learning process (**Fig. 1b**, **Fig. S1b**). In the absence of motion artifacts, *A* is set as an identity operator and excluded from the learnable parameters.

The image-formation model consists of three components: point spread function (PSF, *h*), structure (*s*), and baseline (*b*) (**Fig. 1c**). The PSF *h* is computed as^15^:

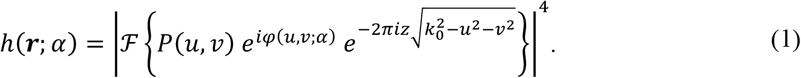

Here, ***r*** represents the spatial coordinates (*x, y, z*) near the focal plane. *P*(*u, v*) and *φ*(*u, v*; *α*) stand for the amplitude and phase maps in the coordinates (*u, v*) within the circular pupil of the objective lens, respectively. *φ*(*u, v*; *α*) is a linear combination of Zernike modes with their associated coefficients *α*, with 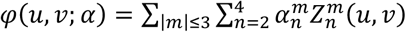. Here *m* and *n* stand for the angular meridional frequency and radial order, respectively, following the American National Standards Institute (ANSI) standard convention for Zernike modes. *α*, a 1D tensor, is a set of learned Zernike coefficients (**Fig. S1c**). We only consider 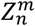 with 2 ≤ *n* ≤ 4 and |*m*| ≤ 3, resulting in 10 Zernike modes to be learned, as including higher-order modes leads to inaccurate aberration estimation.

3D structure *s* is rendered by a neural field (**Fig. 1c**, **Fig. S1a**). It receives the spatial coordinates ***r*** as input and involves both spatial encoding in the Fourier domain^16,17^ and a multilayer perceptron^13,18^. *s* is parametrized by the network weights *θ* and referred to as *f*_*θ*_(***r***).

The baseline term *b*(***r***) is modeled as the multiplication of three 2D tensors that represent baseline elements along each of the *x, y*, and *z* axes (**Fig. 1c**, **Fig. S1d**). This term accounts for both the offset due to background fluorescence and noise and, if present, signal decrease along the *z* axis due to scattering and absorption by tissue.

The image-formation model computes an image stack *ĝ* from PSF *h*(***r***; *α*), structure *s* = *f*_*θ*_ (***r***), and baseline *b*(***r***) by convolving PSF with structure before summation with baseline:

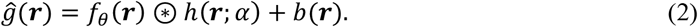

NeAT then compares the input stack *g* (or more generally with motion correction, *Ag*) and the computed stack *ĝ*. It runs an optimization process to update the learnable parameters over iterations to minimize the loss function (**Fig. 1c**, **Fig. S1e**):

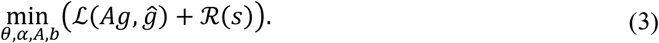

The fidelity term ℒ (*Ag, ĝ*) is a convex combination of SSIM (Structural Similarity Index Metric)^19^ and rMSE (relative Mean-Squared Error)^20–22^ between the two stacks, where SSIM measures the similarity between *Ag* and *ĝ* and rMSE computes a weighted L2 loss that reduces the influence of bright pixels and places greater emphasis on minimizing errors in dark regions. The regularization term ℛ(*s*) incorporates a generic prior on the spatial piecewise smoothness of the structure and is the summation of three regularizations based on second-order total variation^23,24^, L1, and nonlinear diffusion^25^. Second-order total variation and L1 regularizations are chosen for rendering spatially sparse structural features (e.g., sparsely labeled neurons). Nonlinear diffusion regularization is employed to avoid both low-frequency and high-frequency artifacts in the structure recovered by NeAT. More detailed information about the image-formation model, loss function, regularization, and two-step learning is available in **Methods**.

### Performance validation with DWS-AO

To evaluate the accuracy of NeAT’s aberration estimation, we compared the aberration output by NeAT with the ground-truth aberration from DWS with a Shack-Hartmann wavefront sensor of fluorescence from 2P-excited guide stars^26,27^, using a custom-built 2P microscope with perfect conjugation between optics, including between the X and Y galvos (**Fig. S2a**). Microscope system aberration was measured with DWS and corrected by a DM prior to all experiments.

We first validated NeAT’s performance using 2P imaging of fixed Thy1-GFP line M mouse brain slices. A #1.5 coverslip was placed above a brain slice at a 3° tilt, which introduced aberrations similar to those typically induced by a cranial window during *in vivo* mouse brain imaging^28^. We set the correction collar of the objective lens to 0.17, the nominal thickness of the coverslip. From an input image stack (**Fig. 2a**), NeAT output 3D neuronal structures whose lateral (*xy*) and axial (*xz*) maximal intensity projections (MIPs) showed neuronal processes as well as synaptic structures such as boutons and dendritic spines (**Fig. 2b**). The estimated aberration had a similar phase map to the DWS measurement with a root mean square (RMS) difference of 0.09 wave (**Fig. 2c**) and comparable coefficients in the dominant aberration modes, e.g., primary coma 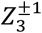 (**Fig. 2d**).

**Figure 2.**
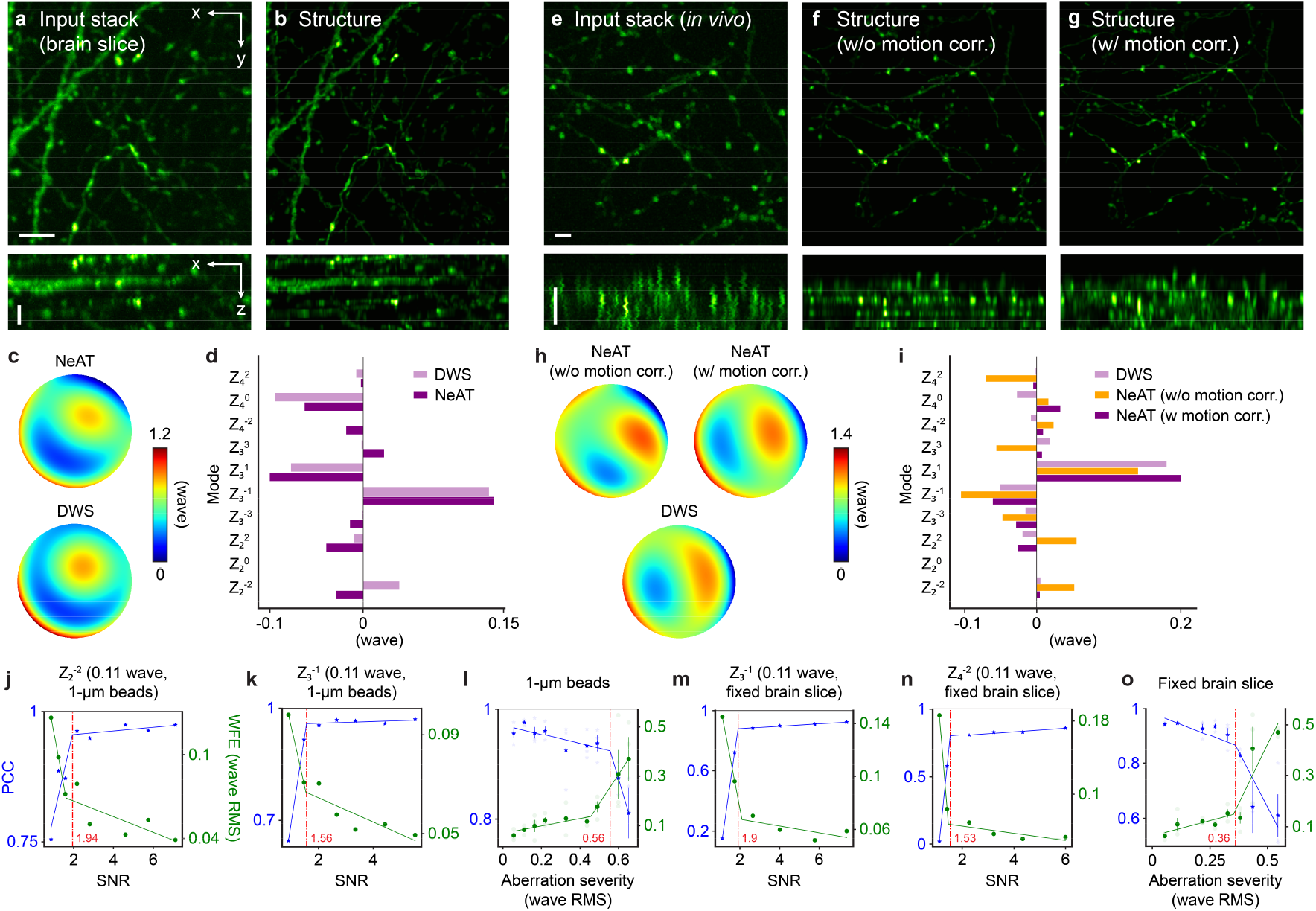
Performance characterization with direct wavefront sensing (DWS) AO in a custom-built 2P microscope. **(a)** Lateral (*xy*) and axial (*xz*) maximal intensity projections (MIPs) of an input image stack to NeAT from a fixed Thy1-GFP line M mouse brain slice. **(b)** Lateral and axial MIPs of the 3D neuronal structure recovered and **(c)** aberration estimated by NeAT, as well as aberration measured by DWS. **(d)** Zernike coefficients of aberrations in **c**. **(e)** Lateral and axial MIPs of an input image stack acquired *in vivo* from a Thy1-GFP line M mouse brain, with motion artifacts visible in *xz*. **(f**,**g)** Lateral axial MIPs of the structures recovered by NeAT without **(f)** and with **(g)** motion correction. **(h)** Aberrations estimated by NeAT without and with motion correction, respectively, and measured by DWS. **(i)** Zernike coefficients for aberrations in **h. (j,k)** Performance versus SNR using 1-µm-diameter beads under primary astigmatism **(j)** and primary coma **(k).** PCC: Pearson correlation coefficient between recovered structures; WFE: wavefront error. **(l)** Performance versus aberration severity evaluated using 1-µm-diameter beads. **(m**,**n)** Performance versus SNR using a fixed mouse brain slice under primary coma **(m)** and secondary astigmatism **(n). (o)** Performance versus aberration severity evaluated using brain slice. Red dashed lines: Cutoff SNRs (**j**,**k**,**m**,**n**) and cutoff aberration RMS (**l**,**o**) from piecewise linear fits (green and blue lines). **(l**,**o)** Data are presented as mean values +/-s.e.m. (N = 3). Scale bars: 5 µm.

Next, we applied NeAT to *in vivo* 2P imaging of the mouse cortex. In one mouse, breathing caused lateral shifts between images at different *z* (**Fig. 2e**). Without correcting for these motion artifacts, the algorithm misinterpreted the laterally displaced images of the same structure at different *z* as separate structures, leading to striated appearance in the axial MIP of its structural output (**Fig. 2f**). NeAT addressed this by using affine transformations *A* to register the image stack, with the transformation matrices jointly learned alongside other parameters (**Eq. 3**). With sample motion corrected, the structural output was free of striation artifacts (**Fig. 2g**), and the aberration output much more closely resembled the ground truth (an RMS error of 0.07 wave) than the output without motion correction (an RMS error of 0.16 wave) (**Fig. 2h,i**).

The effectiveness of sample motion correction depends on the SNR of fluorescence images (**Methods**) and the magnitude of sample motion (**Fig. S3**). For high SNR images (e.g., SNR of 12), NeAT could handle sample motions of ±2 µm of maximum displacement. For noisier images (e.g., SNR of 3), its accuracy decreased and could only handle sample motions with < 0.5 µm displacement. This finding offers practical guidance for optimizing surgical preparation or controlling anesthesia level to minimize sample motion during image acquisition for AO, particularly during deep tissue imaging when SNR is low.

### Performance limit characterizations

After validating NeAT’s performance both *in vitro* and *in vivo*, we evaluated how robustly it performed at varying SNR levels. We varied post-objective power and acquired image stacks of 1-µm-diameter fluorescence beads at different SNRs with primary astigmatism (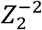, **Fig. S4a**) or primary coma (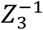, **Fig. S4b**) introduced to the DM. At low SNR levels (e.g., SNR < 1.5), fluorescent beads were poorly visualized and the structures output by NeAT appeared fragmented as they were fitted to noise. Only at sufficiently high SNRs did the structure resemble beads (**Fig. S4a, b**). We quantitatively evaluated NeAT’s performance to identify the cutoff SNR below which NeAT’s performance deteriorated abruptly^29^. We computed the Pearson correlation coefficient (PCC) between the recovered structures at different SNRs and that from an image stack acquired with no aberration and high SNR (SNR > 7, **Fig. S4a, b**). By fitting the PCC values to a piecewise linear curve with two distinct slopes, we identified the cutoff SNR as 1.94 for astigmatism (**Fig. 2j**) and 1.56 for coma (**Fig. 2k**). Below the cutoff SNRs, the accuracy of structural recovery decreased, as indicated by an abrupt drop of PCC values (blue curve, **Fig. 2j,k**); The accuracy of aberration estimation also decreased, as indicated by an increase in wavefront error (quantified by the RMS error between NeAT’s estimate and ground-truth aberrations; green curve, **Fig. 2j,k**).

We repeated the experiment on a fixed Thy1-GFP line M mouse brain slice to determine whether similar limits applied to spatially extended biological structures. In this case, we applied primary coma (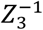, **Fig. S4c**) and secondary astigmatism (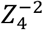, **Fig. S4d**) to the DM separately. Similarly to beads, low-SNR images were associated with structures dominated by artifacts (**Fig. S4c, d**). As before, we calculated the PCC between the recovered structures at different SNRs and the ground truth from an image stack acquired with no aberration and high SNR (SNR > 5, **Fig. S4c, d**). We found that the cutoff SNR was 1.90 for coma (**Fig. 2m**) and 1.53 for astigmatism (**Fig. 2n**), similar to the cutoff SNRs from the bead data. This suggests that at sufficiently high SNRs (SNR ≳ 3 for aberrations tested here), NeAT achieves accurate structural recovery, independent of feature characteristics.

Moreover, we characterized NeAT’s performance limit in terms of aberration severity. We randomly generated Zernike coefficients to obtain mixed-mode aberrations with RMS values ranging from 0.05 to 0.65 waves. We then applied each aberration to the DM and acquired images of beads and brain slices at SNR > 8. With the increase in aberration, fluorescence images became more degraded in resolution and contrast (**Fig. S5**). At the largest aberrations tested (e.g., 0.65 waves for beads and 0.43 waves for brain slices), the recovered structures no longer accurately represented the features of the beads or neurons. We computed the PCC between the structures retrieved by NeAT from images with varying levels of external aberration and the structure from an image stack without aberration. Similar to above, we defined the cutoff RMS as the value above which the PCC exhibited a sudden drop, as identified by fitting the PCC values to a piecewise linear curve with two distinct slopes. We found a cutoff RMS of 0.56 wave for 1-µm beads (**Fig. 2n**) and 0.36 wave for the brain slice (**Fig. 2o**), respectively. This difference in cutoff RMS values is expected as 3D extended structures generally pose greater challenges than beads.

Lastly, we characterized NeAT’s performance limit in terms of sampling rate by varying the pixel sizes of input image stacks. We downsampled both *in vitro* and *in vivo* image stacks of neurons by different factors to vary the input pixel size along the lateral (*dx, dy*) and axial (*dz*) axes, and compared NeAT’s performance in structural recovery and aberration estimation (**Fig. S6, 7**). When pixel size exceeded the Nyquist sampling criterion, the structure outputs from NeAT became inaccurate. The aberration estimation also deviated from the ground truth measured by DWS, with the estimated aberration matching the DWS measurement until lateral pixel size exceeded 0.20 µm and axial pixel size exceeded 0.67 µm, values dictated by the Nyquist condition, for both *in vitro* (**Fig. S6a, Fig. S7a**) and *in vivo* (**Fig. S6b, Fig. S7a**) cases.

### NeAT corrects for conjugation errors in a commercial microscope

Having demonstrated the successful application of NeAT in a custom-built 2P fluorescence microscope and acquired a thorough understanding of its performance in relation to SNR, motion, aberration severity, and input pixel size, we next tested whether NeAT worked on a commercial 2P microscope. This was motivated by the desire to expand the application of AO beyond optical specialists to a general laboratory setting with microscopes having imperfect conjugation and misalignment of optical components, as well as limited access and adjustability of their optical paths.

We added a liquid-crystal SLM to the beam path between an excitation laser and a commercial 2P fluorescence microscope (Bergamo II, Thorlabs) (**Fig. S2b**). This system differs from our custom-built microscope in several ways. First, the DM, *x* galvo, and *y* galvo of the custom-built system were conjugated with pairs of lenses (**Fig. S2a**) to ensure that the corrective phase pattern displayed on the DM was accurately relayed to the back focal plane (BFP) of the objective lens and stayed stationary during beam scanning. But the commercial microscope, as typical for microscopes in biological laboratories, did not conjugate the two galvos but placed them close to each other. Second, while the optics of the custom-built system were carefully arranged and aligned to ensure the registration between the *x* and *y* axes of the SLM surface and the fluorescence images, the commercial microscope had multiple mirrors in an enclosed optical path whose placement and alignment were preset and not adjustable. Finally, the commercial system was designed to have the whole microscope body move in 3D to accommodate large samples, which causes alignment errors between the SLM on the optical table and the objective lens in the microscope that for heavily shared microscopes can vary daily. As a result, a wavefront applied to SLM is translated, rotated, scaled, and/or sheared at the BFP of the objective lens of the commercial microscope, which in turn degrades the performance of aberration correction.

To address this problem, we incorporated into NeAT a process to estimate and correct conjugation errors of commercial microscopes (**Fig. 1a**). Corrective wavefront displayed on the SLM, *φ*_Corr_, becomes *φ*_BFP_ at the BFP of the objective lens, with

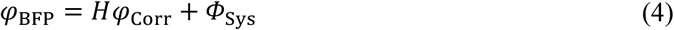

Here *Φ*_Sys_ represents the system aberration and *H* is a linear geometric transformation describing the effects of conjugation errors on *φ*_Corr_ (**Fig. 3a**). We model *H* as an affine transformation with parameters for translational, rotational, scaling, and shear transformation (**Fig. 3b**). For microscopes with perfect conjugation, *H* = *I*, the identity operator (*i*.*e*. translations are 0 pixels in *x* and *y*, rotation is 0 deg, scaling is 1, and shear is 0). For microscopes with conjugation errors, the process of accounting for them requires finding the transformation *H* and system aberration *Φ*_Sys_.

**Figure 3.**
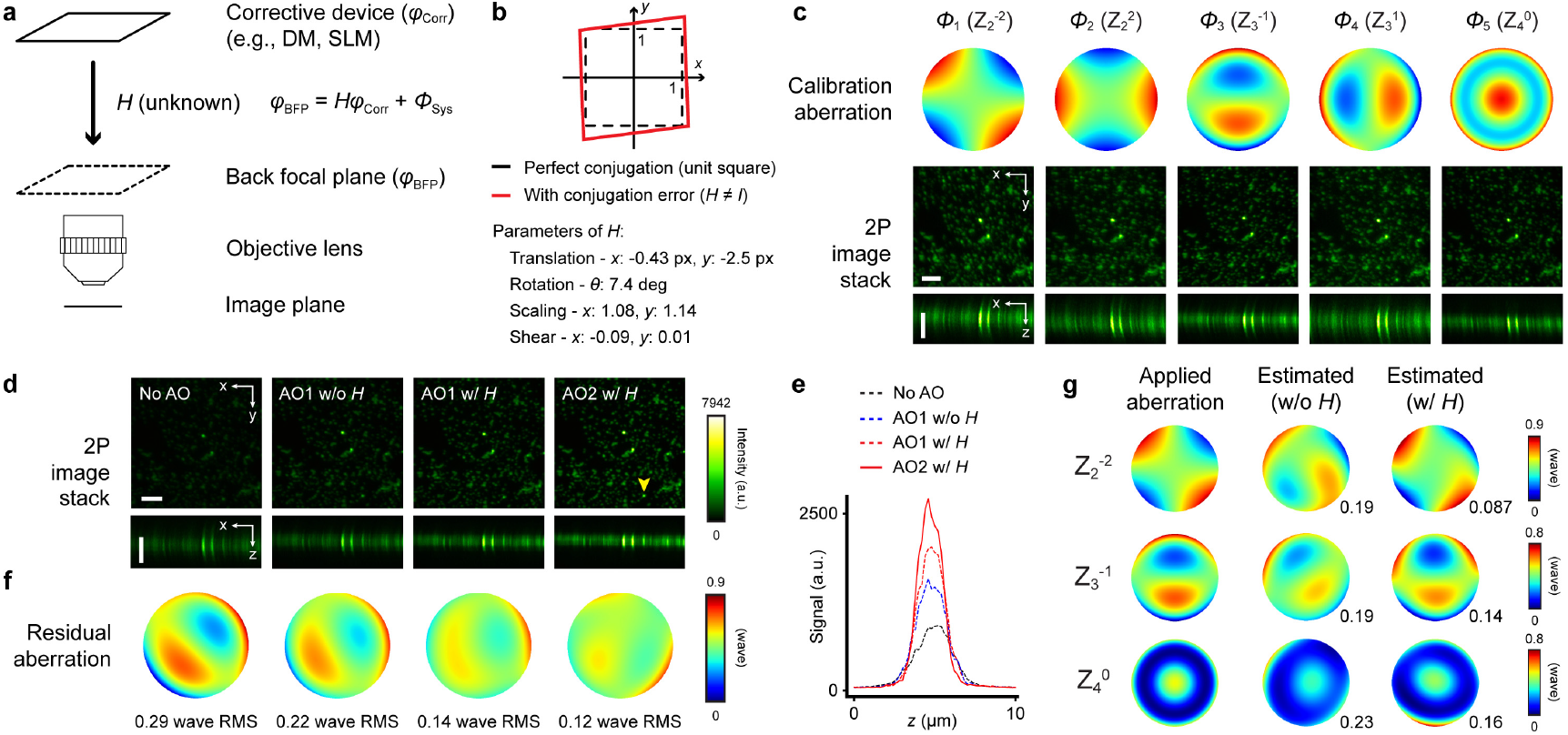
NeAT corrects for conjugation errors in a commercial microscope. **(a)** Conjugation errors transform corrective pattern *φ*_Corr_ on SLM to *φ*_BFP_ = *Hφ*_Corr_ + *Φ*_Sys_ at objective lens back focal plane. *Φ*_Sys_: system aberration. **(b)** *H* (with example affine parameters) translates, rotates, scales, and shears a unit square (black dashed square) to a parallelogram (red). px: pixel. **(c)** *H* is determined from image stacks of 200-nm-diameter beads acquired with calibration aberration *Φ*_*n*_ (*n* = 1,2, ⋯, 5) applied to SLM. Lateral (*xy*) and axial (*xz*) MIPs of the calibration image stacks are shown. **(d)** Lateral and axial MIPs of image stacks of 200-nm-diameter beads acquired without system aberration correction (‘No AO’), after one iteration of AO without (‘AO1 w/o *H*’) or with (‘AO1 w/*H*’) conjugation correction, and after two iterations of AO with conjugation correction (‘AO2 w/*H*’). **(e)** Axial signal profiles of the bead marked by yellow arrowhead in **d. (f)** Residual aberrations estimated by NeAT from image stacks in **d. (g)** Left to right: aberrations (with 0.3 wave RMS) applied to SLM, estimated aberration by NeAT without conjugation correction, and estimated aberration by NeAT with conjugation correction from bead image stacks acquired with the applied aberration. Numbers to the bottom right of estimated aberrations: difference (in wave RMS) between estimated aberration and applied aberration. Scale bars: 5 µm.

We determine system aberration *Φ*_Sys_ by inputting into NeAT an image stack of 200-nm-diameter fluorescence beads acquired with a flat phase pattern applied to the SLM. The estimated system aberration from NeAT is 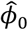, with

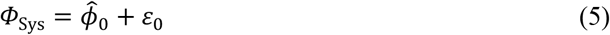

Here *ε*_0_ represents estimation error by NeAT, which should be much smaller in RMS magnitude than *Φ*_Sys_.

To determine *H*, we apply 5 calibration aberrations *Φ*_*n*_ (*n* = 1 to 5) including primary astigmatism (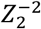 and 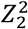), coma (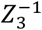 and 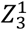), and spherical aberration 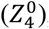, to the SLM. These calibration aberrations allow us to detect translation, scaling, rotation, and shear errors in conjugation. At the objective lens BFP, these aberrations became *HΦ*_*n*_ + *Φ*_Sys_. With image stacks of 200-nm fluorescence beads acquired under these external aberrations as inputs (**Fig. 3c**), NeAT Returns 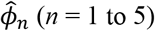, with

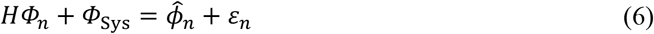

Here *ε*_*n*_ represents estimation error by NeAT. Subtracting (5) from (6) and assuming *ε*_*n*_ – *ε*_0_ ≈ 0, we have

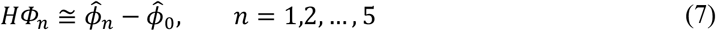

Now with *Φ*_*n*_ (*n* = 1 to 5) known, and 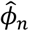 and 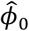 from NeAT, we determine the parameters of *H* by minimizing the loss function:

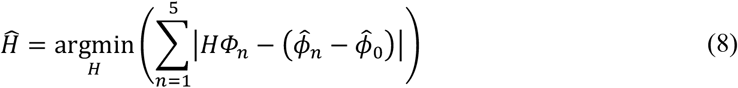

*Ĥ*, the estimate for *H*, describes how the conjugation errors in the system translate, rotate, scale, and shear the wavefront pattern applied to the SLM on its way to the BFP of the objective lens. To correct these errors, we then apply the inverse of *Ĥ, Ĥ*^−1^, to the aberration estimation 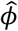 from NeAT and use 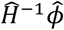 as the corrective pattern on the SLM (**Fig. 1d**).

For example, to correct for system aberration of the commercial microscope, we used an image stack of 200-nm fluorescence beads as input to NeAT, which returned 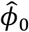 as the aberration estimation. Directly applying 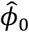 to the SLM increased the signal of a fluorescent bead by 1.7-fold (“AO1, w/o H”, **Fig. 3d,e**). By also correcting for conjugation errors, 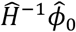 increased the signal by 2.2-fold (“AO1, w/ H”, **Fig. 3d,e**). Using the image stack acquired with 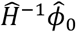 as input into NeAT, we obtained the residual aberration 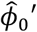 and applied 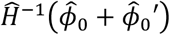 to the SLM, leading to a 3.0-fold signal gain over no aberration correction (“AO2, w/ H”, **Fig. 3d,e**). From the image stacks acquired with these corrective patterns, NeAT estimated the residual aberrations (**Fig. 3f**). Consistent with the fluorescent signal measurements, conjugation error correction substantially reduced residual aberration, with 0.14 and 0.12 wave RMS after the first and second iterations of AO correction, while the residual aberration without conjugation correction had a 0.22 wave RMS.

We further tested our approach on correcting known astigmatism, coma, and spherical aberrations introduced to the SLM. From bead image stacks acquired with these aberrations applied, NeAT returned estimated aberrations (“Estimated w/o H”, **Fig. 3g**), which represented the wavefront distortion at the objective BPF and substantially differed from the applied aberrations (“Applied aberration”, **Fig. 3g**) due to conjugation errors. Transforming the estimated aberration with *Ĥ*^−1^, we obtained aberrations with phase maps (“Estimated w/ H”, **Fig. 3g**) that closely matched the given aberrations in all three cases, leading to much smaller RMS errors (astigmatism: 0.087 and 0.19 wave RMS with and without *H* correction; coma: 0.14 and 0.19 wave RMS with and without *H* correction; spherical: 0.16 and 0.23 wave RMS with and without *H* correction). Once characterized, the same *Ĥ*^−1^ can be applied as long as the conjugation of the microscope remains unchanged. Below, the system aberration of the commercial microscope was always corrected for “No AO” images so that improvement by AO arose from the correction of sample-induced aberrations alone.

### Real-time aberration correction for *in vivo* structural imaging of mouse brain

We evaluated NeAT’s capacity to improve *in vivo* structural imaging with the commercial microscope. We acquired an image stack of a tdTomato-expressing dendrite at 350 µm depth in the primary visual cortex (V1) of a head-fixed mouse (“No AO”, **Fig. 4a**) and used it as input to NeAT. Applying to the SLM the corrective wavefront from NeAT with both motion and conjugation corrections, we imaged the same dendrite and observed a marked improvement in brightness (up to 1.8× for dendritic spines), resolution, and contrast (“Full correction”, **Fig. 4a**).

**Figure 4.**
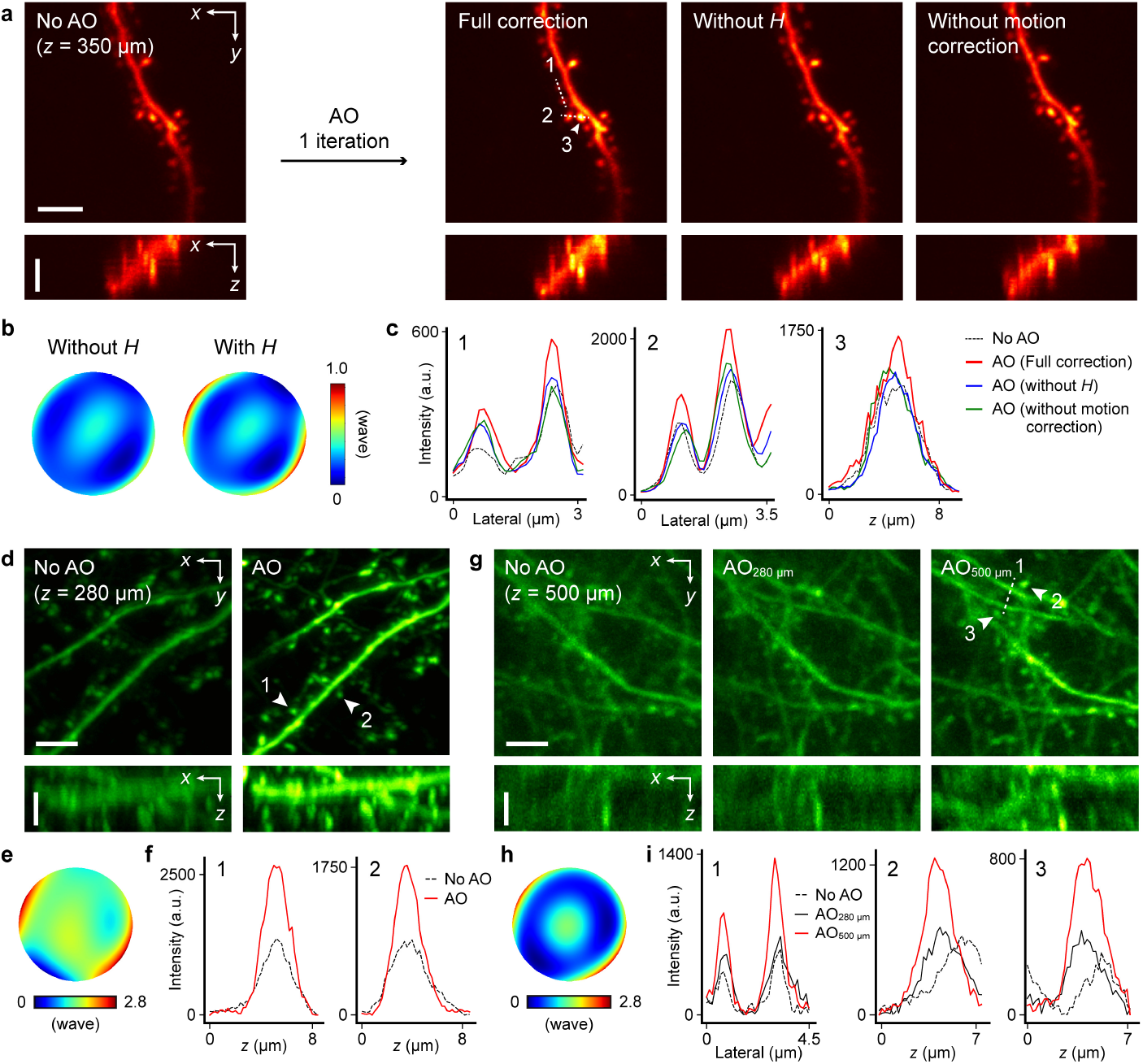
Real-time aberration correction by NeAT for *in vivo* structural imaging. **(a)** Lateral (*xy*) and axial (*xz*) MIPs of image stacks of tdTomato-expressing dendrite and dendritic spines at 350 µm depth acquired with system aberration correction only (“No AO”, used as input to NeAT), with corrective wavefront estimated by NeAT with both conjugation and motion corrections (“Full correction”), motion correction only (“Without H”), or conjugation correction only (“Without motion correction”). **(b)** Estimated aberrations by NeAT without and with conjugation correction. **(c)** Lateral signal profiles along dashed lines and axial signal profiles of spine indicated by arrowhead in **a. (d)** Lateral and axial MIPs of Thy1-GFP line M mouse dendrites at 280 µm depth acquired with system aberration only (“No AO”, used as input to NeAT) and with correcting sample-induced aberration by NeAT. **(e)** Estimated aberration by NeAT. **(f)** Axial profiles of dendritic spines marked by arrowheads in **d. (g)** Lateral and axial MIPs of neuronal processes at 500 µm depth, acquired with system aberration correction only (“No AO”), aberration correction at 280 µm (“AO_280 µm_”, used as input to NeAT; wavefront in **e**), and aberration correction at 500 µm (“AO_500 µm_”) **(h)** Sample-induced aberration at 500 µm. **(i)** Lateral profiles along dashed line and axial profiles of spines indicated by arrowheads in **g**.Scale bars: 5 µm.

Correcting for both sample motion and conjugation error was necessary for the observed improvement. The corrective wavefront with only motion correction but not conjugation correction substantially differed from that with full correction (**Fig. 4b**) and led to more modest improvements in image quality (“Without *H*”, **Fig. 4a**). Only correcting for conjugation but not motion similarly underperformed (“Without motion correction”, **Fig. 4a**). These trends were observed quantitatively in the lateral and axial intensity profiles of three example dendritic spines (**Fig. 4c**).

We investigated further whether image-registration software such as the StackReg plugin in ImageJ can work similarly well to the motion correction method integrated into the learning process of NeAT. We acquired an image stack of beads with aberration, introduced simulated motion artifacts, pre-registered the stack with StackReg, and then used the resulting image stack as input to NeAT. Although structural recovery was moderately successful for beads (**Fig. S8a, b**), the accuracy of aberration estimation from the pre-registered image stack was inferior to inputting the un-registered stack to NeAT directly (**Fig. S8c**). Similarly, pre-registration with StackReg on an image stack acquired *in vivo* led to a corrective wavefront with smaller brightness improvement than the corrective wavefront from motion correction by NeAT (**Fig. S8d**,**e**). This can be explained by whether motion correction considers the existence of aberration. While NeAT integrates motion correction into its learning process for aberration (**Eq. 3**), conventional image registration is unaware of aberrations and matches features between adjacent image planes to align them, which may inadvertently reduce or exaggerate certain aberrations (e.g., the axially curved tail of comatic aberration may be straightened by StackReg).

Having established that both conjugation and motion corrections are needed for *in vivo* imaging using the commercial microscope, we further tested NeAT’s performance for morphological imaging deep within the brain of a Thy1-GFP line M mouse. We first used an image stack acquired at a depth of 280 µm as input to NeAT (“No AO”, **Fig. 4d**) to obtain the corrective wavefront (**Fig. 4e**, 0.36 wave RMS,), which led to resolution improvement as well as an ∼2× increase in spine brightness (“AO”, **Fig. 4d,f**). We then acquired an image stack at 500 µm depth while applying to the SLM the corrective wavefront at 280 µm (“AO_280 µm_”, **Fig. 4g**). Using the image stack as input to NeAT, we obtained a corrective wavefront, which was then added to the corrective wavefront at 280 µm to obtain the final corrective pattern (**Fig. 4h**, 0.49 wave RMS). This corrective wavefront has a larger RMS magnitude than that at 280 µm, consistent with previous observation of stronger aberrations at larger imaging depths for the mouse brain^30^. Compared to the image stacks acquired without AO (“No AO”, **Fig. 4g**) and with corrective wavefront at 280 µm (“AO_280 µm_”, **Fig. 4g**), images after correction at 500 µm (“AO_500 µm_”, **Fig. 4g**) had the best resolution and contrast, with up to a 2.4-fold increase in brightness for dendritic and synaptic structures (**Fig. 4i**). Here by using the corrective wavefront at a shallower depth when acquiring the input image stack at a deeper depth, we overcame the limit on aberration severity and used NeAT to correct large aberrations experienced in deep tissue imaging.

### NeAT improves *in vivo* glutamate imaging from the mouse brain

We next used NeAT with motion and conjugation correction to improve *in vivo* functional imaging in head-fixed mice. We expressed the genetically encoded glutamate indicator iGluSnFR3^31^ sparsely in V1 neurons (**Methods**). From an image stack of dendrites at 400-µm depth (**Fig. 5a**), NeAT returned a corrective wavefront (**Fig. 5b**) that substantially increased image resolution and contrast, resulting in approximately two-fold improvement in brightness as shown by axial profiles of dendritic spines (i,ii; **Fig. 5c**) and resolving a dendritic spine from its nearby dendrite (iii; **Fig. 5c**).

**Figure 5.**
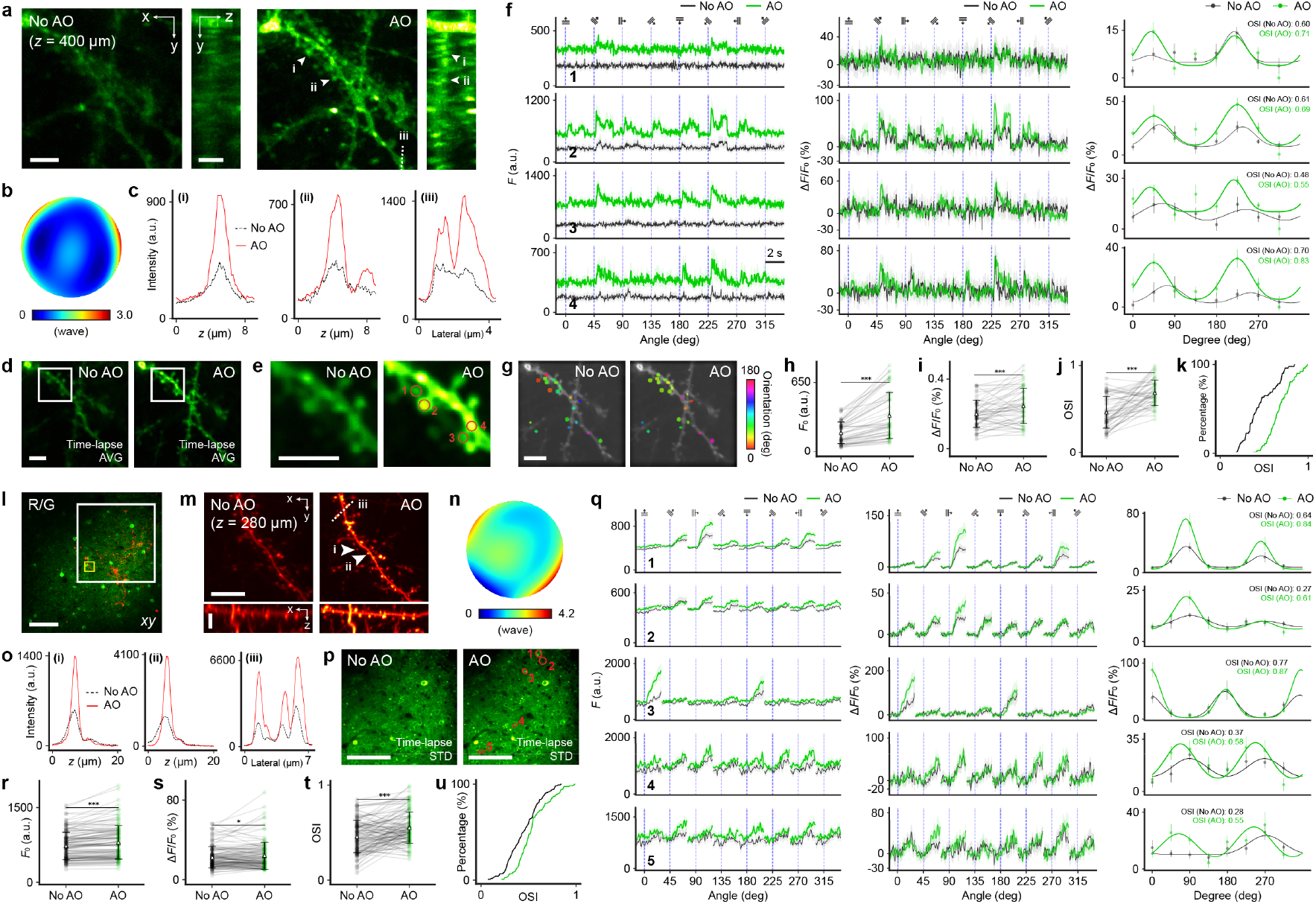
Real-time aberration correction by NeAT for *in vivo* glutamate and calcium imaging. **(a)** Lateral (*xy*) and axial (*yz*) MIPs of input stacks to NeAT (“No AO”) and stacks acquired after aberration correction by NeAT (“AO”) of dendrites expressing iGluSnFR3 at 400 µm depth in mouse V1. **(b)** Estimated aberration by NeAT. **(c)** Axial profiles of spines indicated by arrowheads and lateral profiles along dashed line in **a. (d)** Averages of time-lapse *xy* images of dendrites measured without and with AO. **(e)** Zoomed-in views of structures in box in **d. (f)** Trial-averaged signal traces (*F*), glutamate transient traces (Δ*F/F*_0_), orientation tuning curves and OSI values of 4 ROIs (1-4 in **e**). Shade and error bars: s.e.m. **(g)** OS spines in **d** color-coded by their preferred orientations measured without and with AO. **(h-k)** Comparisons of basal fluorescence (F_0_; **h**), glutamate transient (Δ*F/F*_0_; **i**), OSIs (**j**,**k**) of 52 OS spines out of 86 total spines before and after AO correction. Two-sided paired t-test (**h-j**), *p* < 0.001. Kolmogorov-Smirnov test (**k**), *p* < 0.001. **(l)** Superimposed *xy* images of sparse tdTomato-expressing neurons (1000-nm excitation) and dense GCaMP6s-expressing neurons (920-nm excitation) at 280 µm depth. **(m)** Lateral (*xy*) and axial (*xz*) MIPs of dendrites (yellow box in **l**) measured without (“No AO”, input to NeAT) and with AO. **(n)** Estimated aberration by NeAT. **(o)** Axial profiles for spines indicated by arrowheads and lateral profiles along dashed line in **m. (p)** Standard deviation of time-lapse images of GCaMP6s-expressing neurons in white box in **l**, acquired without and with AO. **(q)** Trial-averaged signal traces (*F*), calcium transient traces (Δ*F/F*_0_), orientation tuning curves, and OSI values of 5 ROIs (1-5 in **p**). Shade and error bars: s.e.m. **(r-u)** Comparisons of basal fluorescence (F_0_; **h**), calcium transient (Δ*F/F*_0_; **s**), OSIs (**t**,**u**) of 125 OS ROIs out of 255 somatic and neuronal structures before and after AO correction. Two-sided paired t-test, *p* < 0.001 (**r**,**t**), *p* < 0.05 (**s**). Kolmogorov-Smirnov test (**u**), *p* < 0.001. Scale bars: (**a**,**d**,**e**,**g**) 5 µm; (**l**,**p**) 100 µm; (**m**) 10 µm.

Subsequently, we presented gratings drifting in eight different directions (0°, 45°, ⋯, 315°; 10 repetitions) to the mouse and recorded time-lapse images of dendritic spines in the same FOV as in **Fig. 5a**, with and without aberration correction at a 60 Hz frame rate. With iGluSnFR3 labeling, changes in fluorescence brightness reflected glutamate release and thus synaptic input strength at these dendritic spines. Consistent with the above result, AO increased the brightness of dendrites and spines in the averaged time-lapse image (**Fig. 5d**; **Fig. 5e**, zoomed-in views of white boxes in **Fig. 5d**). For four representative dendritic spines (ROI 1-4, **Fig. 5e**), AO correction doubled the basal intensity (*F*_0_) of their trial-averaged fluorescent traces and led to more prominent glutamate transients with larger amplitudes (Δ*F/F*_0_) (left and middle panels, **Fig. 5f**). Fitting the glutamate responses to the 8 drifting grating stimuli with a bimodal Gaussian curve^32^, we obtained the orientation-tuning curves for these spines (right panels, **Fig. 5f**). Here AO increased the response amplitudes to the preferred grating orientations and led to a higher orientation sensitivity index (OSI) for these spines. Correcting aberration also shifted the preferred orientation of some spines (e.g., ROI 3 and 4, **Fig. 5f**), resulting in more similar tuning preference for neighboring spines (**Fig. 5g**), consistent with previous findings^33^. Consistently across spine populations (52 orientation-sensitive ROIs out of 86 spines, **Methods**), aberration correction by NeAT significantly increased basal fluorescence F_0_ by 1.9-fold on average (two-sided paired t-test, *p* < 0.001, **Fig. 5h**). It also increased Δ*F/F*_0_ and OSI values as indicated by pairwise comparison (two-sided paired t-test, *p* < 0.001, **Fig. 5i** and **Fig. 5j**, respectively) and the cumulative OSI distributions (Kolmogorov-Smirnov test, *p* < 0.001, **Fig. 5k**).

### NeAT improves *in vivo* calcium imaging in densely labeled brains

We further demonstrated that NeAT can also be applied to densely labeled brains, a common application scenario for *in vivo* calcium imaging of neuronal populations. As NeAT requires an input stack of sparse structures for aberration estimation, we used viral transduction to densely express the genetically encoded calcium indicator GCaMP6s^34^ and sparsely express the red fluorescent protein tdTomato in L2/3 neurons of the mouse V1 (**Methods**). Because aberration estimation and correction can be performed at different excitation wavelengths without compromising correction performance (**Fig. S9**), we used an image stack of a tdTomato-expressing neuron (inside yellow box of **Fig. 5l**) acquired with 1000 nm excitation light as the input to NeAT (32×32×10 µm^3^ stack, “No AO”, **Fig. 5m**) to obtain the corrective wavefront (**Fig. 5n**). AO visibly improved image contrast and resolution of the tdTomato-expressing neuron (“AO”, **Fig. 5m**), leading to a >2× increase in intensity in both axial profiles at dendritic spines and lateral profiles across dendrites (**Fig. 5o**).

Next, we switched the excitation wavelength to 920 nm and acquired images of GCaMP6s-expressing neurons over a 484×484 µm^2^ FOV (green channel of **Fig. 5l**) without and with the corrective wavefront obtained by NeAT at 15 Hz, while presenting drifting gratings to the head-fixed mouse to evoke calcium responses. The standard deviation images of the time-lapse stacks showed greater intensity differences across time frames after aberration correction (**Fig. 5p**, zoomed-in views on the white boxes in **Fig. 5l**), indicative of larger calcium transient magnitude. Indeed, for five representative ROIs (1-5, **Fig. 5p**), calcium transients were more apparent and had larger magnitudes in both trial-averaged fluorescence (*F*) and Δ*F/F*_0_ traces with AO (left and middle panels, **Fig. 5q**), leading to higher orientation selectivity indices for these structures (right panels, **Fig. 5q**).

Over the population of 125 orientation-selective ROIs out of 255 somatic and neuronal structures within the whole FOV, we found statistically significant differences between No AO and AO conditions for both basal fluorescence F_0_ (two-sided paired t-test, *p* < 0.001, **Fig. 5r**) and Δ*F/F*_0_ (*p* < 0.05, **Fig. 5s**). Here the increase in basal fluorescence was less than what we observed for glutamate imaging of dendritic spines, because aberration decreases signal brightness of smaller structures such as dendritic spines more than larger structures such as somata^30,35,36^. Similar to glutamate imaging, AO increased the OSIs of neuronal structures (two-sided paired t-test, *p* < 0.001, **Fig. 5t**; for cumulative distributions of OSI, Kolmogorov-Smirnov test, *p* < 0.001, **Fig. 5u**).

## Discussion

In this work, we describe NeAT, a general-purpose AO framework for aberration measurement and correction for 2P microscopy using neural fields. Neural fields refer to implicit functions represented by a coordinate-based neural network across spatial coordinates^14^. They have been used for various computational imaging applications^16,29,37–41^, including Neural Radiance Fields (NeRF)^13^ for 3D scene representation. NeAT has several distinct features that set it apart from NeRF and other neural field applications (detailed in **Table S1**), including its incorporation of a physics-based prior specific to the 2P imaging system, its estimation and correction of sample motion and microscope conjugation errors, and its joint recovery of 3D structural information alongside aberration estimation.

Using the physics-based prior of the 2P imaging system, implemented through an image-formation model that accounts for both aberrations and sample motion, NeAT accurately estimates optical aberrations from a 2P fluorescence image stack without the need for external supervision, even in the presence of motion artifacts during live animal imaging. Importantly, this functionality eliminates the need for integrating AO capabilities into microscope control software, making it applicable to existing custom-built and commercial 2P microscopy systems in general.

Furthermore, NeAT can estimate and correct conjugation errors. Such errors are common in homebuilt and commercial microscopes used in general biology laboratories and cause the distortion of the applied corrective phase pattern at the objective lens back focal plane, leading to deteriorated AO performance. NeAT measures the impact of conjugation errors on calibration aberrations and compensates for them by preemptively transforming the corrective phase pattern before it is displayed on the wavefront-shaping device. This feature would greatly facilitate the broader adoption of AO in a wide range of microscope systems.

Additionally, NeAT simultaneously recovers 3D structural information while estimating aberration during its learning process. For applications where only structural information is needed, this unique capability eliminates the requirement of wavefront-shaping devices or the need for additional imaging with AO correction, greatly lowering system complexity and cost.

We rigorously evaluated NeAT’s performance under various conditions, such as different SNR levels, aberration severity, and motion artifacts. We established the framework’s performance limits for accurate and reliable operation, providing valuable guidelines on imaging settings when applying NeAT.

Finally, we applied NeAT to *in vivo* structural and functional imaging of the mouse brain by a commercial microscope, demonstrating its capability in improving image quality for demanding real-life biological applications. NeAT effectively estimates and corrects aberrations deep into the mouse brain, enabling morphological imaging of synapses at 500 µm with improved resolution and contrast. It also improves the signal and accuracy of glutamate and calcium imaging of synapses and neurons in the mouse visual cortex responding to visual stimuli.

With a single z-stack as input and a computation time of a few minutes (Table S2), NeAT’s simple implementation, robust performance, and ability to correct for motion and conjugation errors in imaging systems offer great potential for broader adoption and impact in biological research.

## Methods

### Custom-built two-photon microscopy with direct wavefront sensing AO

A custom-built two-photon fluorescence microscope was equipped with a wavefront sensor for DWS and described previously^42,43^ (**Fig. S2a**). A Ti-Sapphire laser (Chameleon Ultra II, Coherent Inc.) was tuned to 920 nm output and scanned by a pair of carefully conjugated galvos (H2105, Cambridge). Pairs of achromatic doublet lenses (L3-L8) conjugated the surfaces of galvos with a DM (PTT489, Iris AO) and the BFP of an objective lens (CFI Apo LWD ×25, 1.1 NA, 2.0 mm WD, Nikon). During imaging, 2P excited fluorescence was collected by the same objective, reflected by a dichroic mirror (D2, Di02-R785-25×36, Semrock), and detected by a GaAsP photomultiplier tube (H7422-40, Hamamatsu). For wavefront sensing, the emitted 2P fluorescence was descanned by the galvo pair, reflected by a dichroic mirror (D1, Di02-R785-25×36, Semrock), and directed to a Shack-Hartmann (SH) sensor through a pair of achromatic lenses (FL = 60, 175 mm). The SH sensor consisted of a lenslet array (Advanced Microoptic Systems GmbH) conjugated to the objective BFP and a CMOS camera (Orca Flash 4.0, Hamamatsu) positioned at the focal plane of the lenslet array.

### Commercial two-photon fluorescence microscopy with an AO module

The commercially available multiphoton microscope (Bergamo II, Thorlabs) used a Ti-Sapphire laser (Chameleon Ultra II, Coherent Inc.) tuned to 920 nm or 1000nm for 2P excitation. An AO module consisted of a liquid crystal SLM (1024 × 1024, HSP1K, Meadowlark Inc.) and two pairs of relay lenses (L1-L4, FL = 200, 50, 500, and 500mm) was added to the beam path on the optical table between the laser and the microscope. The laser output had its polarization rotated by an achromatic half-wave plate (AHWP05M-980, Thorlabs) to align with the SLM polarization requirement and was expanded 15 times using two beam expanders (GBE03-B, GBE05-B, Thorlabs) to fill the active area of the SLM. The two pairs of relay lenses demagnified the laser output and conjugated the SLM surface to the non-resonant galvo surface within the galvo-resonant-galvo scanning head of the microscope. A pair of scan lenses within the Bergamo II microscope (L5-L6, FL=50 and 200mm) relayed the laser to the BFP of a water-dipping objective (25×, 1.05 NA, 2mm WD, Olympus). Fluorescence emission was collected through the objective and detected by two GaAsP photomultiplier tubes (PMT 2100, Thorlabs) for two-color imaging of green (525/50 nm emission filter) and red (607/70 nm emission filter) fluorescence, respectively.

### Animals and surgical procedures

All animal experiments were conducted in accordance with the National Institutes of Health guidelines for animal research. Procedures and protocols involving mice were approved by the Institutional Animal Care and Use Committee at the University of California, Berkeley. *In vivo* imaging experiments were performed using 2-4-month-old wild-type (C57BL/6J) or Thy1-GFP line M mouse lines.

Cranial window and virus injection surgeries were conducted under anesthesia (2% isoflurane in O_2_) following established procedures^33,44^. For *in vivo* glutamate imaging, sparse expression of iGluSnFR3 was achieved in V1 L2/3 by injecting a 1:1 mixture of diluted AAV2/1-Syn-Cre virus (original titer 1.8 × 10^13^ GC/ml, diluted 10,000-fold in phosphate-buffered saline) and AAV-hSyn-FLEX-iGluSnFR3-v857-SGZ at multiple sites 150-250 µm below pia. 25 nl of the virus mixture was injected at each site. For *in vivo* calcium imaging, dense expression of GCaMP6s and sparse expression of tdTomato was achieved in V1 L2/3 by co-injecting a 1:1:1 mixture of diluted AAV2/1-Syn-Cre virus (original titer 1.8 × 10^13^ GC/ml, diluted 1,000-fold in phosphate-buffered saline), AAV2/1-CAG-FLEX-tdTomato (6 × 10^13^ GC/ml), and AAV1-Syn-GCaMP6s-WPRE-SV40 (1 × 10^13^ GC/ml) at multiple sites 150-250 µm below the pia. 25 nl of the virus mixture was injected at each site. A cranial window, made of a glass coverslip (Fisher Scientific, no. 1.5), was embedded in the craniotomy and sealed in place with Vetbond tissue adhesive (3M). A metal head post was attached to the skull using cyanoacrylate glue and dental acrylic. After 3 weeks of expression and 3 days of habituation for head fixation, *in vivo* imaging was conducted in head-fixed mice under anesthesia (1% isoflurane in O_2_) for structural imaging and in lightly anesthetized mice (0.5% isoflurane in O_2_) for functional imaging.

### Loss function and regularization in self-supervised learning process

The fidelity term ℒ (*Ag, ĝ*) in the loss function (**Eq. 3**) is represented as a convex combination of SSIM^19^ and rMSE^20–22^ as follows:

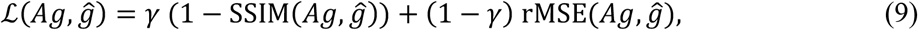

where rMSE is defined as

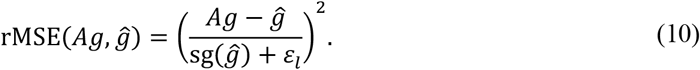

Here sg(·) denotes a stop-gradient operation that treats its argument as a constant, employed for numerical stability during backpropagation^20^. The parameter *γ* controls the weight between the two terms. It is set to 0.25 if the RMS contrast of the image stack’s background pixels, *i*.*e*., *ε*_*l*_ = *σ*_*b*_(*g*_*bfr*_), is larger than 0.03, where *σ*_*b*_(·) computes the standard deviation of the background pixels of the operand. If the contrast is smaller than 0.02, *γ* is set to 1.0. Otherwise, *γ* is linearly interpolated between 0.25 and 1.0. Here, *g*_*bfr*_ represents a background-fluctuation-removed version of *g*, introduced to remove any unwanted low-frequency fluctuations in the images that could otherwise exaggerate the standard deviation.

The regularization term ℛ(*s*) in the loss function (**Eq. 3**) is designed to render spatially sparse and smooth structural details, serving as a generic prior that reflects structural features of mouse brain neurons. It includes three regularization terms: second-order total variation (TV) ℛ_*tv*_(*s*)^23,24^, L1 regularization ℛ_L1_(*s*), and nonlinear diffusion (NLD) ℛ_NLD_(*s*)^25^.

First, second-order TV ℛ_*tv*_(*s*) aims to recover smooth profiles from noisy measurements by sparsifying the spatial gradient components. Unlike first-order TV^45^, which uses first-order derivatives, second-order TV uses second-order derivatives to avoid staircase artifacts^23,24^. In our implementation, we further applied a nonlinear tone mapping function^20^, an approximated logarithmic function (**Eq. 12**) that strongly penalizes errors in regions with low intensity values. For simplicity, the spatial coordinates (*x, y, z*) are expressed as (*x*_1_, *x*_2_, *x*_3_) below.

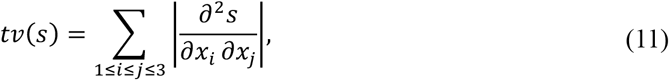

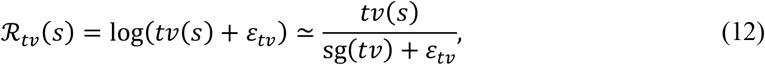

where sg(·) indicates the same stop-gradient operation as above, and *ε*_*tv*_ is determined from the input image stack *g* as the smallest standard deviation of second-order difference *tv*(*g*), that is,

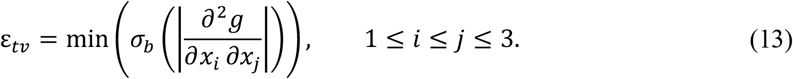

Second, L1 regularization ℛ_L1_(*s*) helps to render the structure *s* with spatially sparse features by adding a penalty based on the absolute value of *s* as follows,

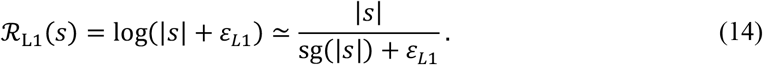

Here *ε*_*L*1_ = *σ*_*b*_(| *g*_*bfr*_|) and the same logarithmic tone mapping function^20^ (**Eq. 12**) is applied on the top of the absolute value.

Lastly, NLD regularization^25^ ℛ_NLD_(*s*) constrains the magnitude of the first-order difference of the structure *s*, computed along the depth axis *z*. This suppresses slowly varying spatial components while preventing the structure from fitting to rapidly varying axial features that sparsity-promoting regularizations might favor. This regularization balances the influence of the first two terms, allowing the structure to retain desirable details. It is written as

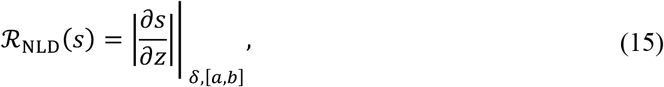

where *f*|_*δ*, [a,b]_ ≡ max(*f, b*) + *δ* max(*a*, min(*f, b*)) + min(*f, a*). For all results presented in this manuscript, *δ* = 0.1, *a* = 0.005, *b* = 2.0.

Together, the summation of the regularization terms is expressed as

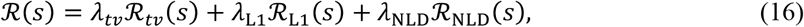

where *λ*_*tv*_ = 0.005, *λ*_L1_ = 0.01, *λ*_NLD_ = 10^δ6^.

### Baseline term *b* in image formation

The baseline term *b* is modeled as low rank to account for the offset due to baseline fluorescence or noise and potential power decrease along the depth axis caused by scattering or absorption in deep tissue imaging. *b* is represented as the sum of rank-1 tensors, and we set to *R* = 5 here:

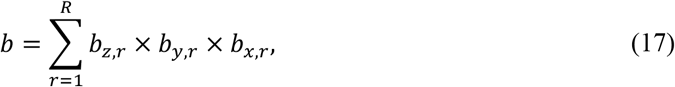

where *b*_*x,r*_, *b*_*y,r*_, *b*_*z,r*_ are learnable 2D tensors to represent baseline components along the *x, y*, and *z* axes, respectively. These tensors are initialized with the value (0.1 *σ*_*b*_(|*g*_*bfr*_|)^1/3^. By limiting the rank of the baseline model, we constrain it to capture only low-spatial-frequency baseline features, thereby effectively separating the baseline from the sample features.

### Two-step learning process

The weights of the neural network *θ* (representing structure *s*), Zernike coefficients *α* (thus PSF *h*(***r***; *α*)), and baseline term *b* in the image-formation model are optimized in a two-step learning process^29^. The first step only adjusts neural network weights for *s*, while the Zernike coefficients *α* and baseline *b* remain fixed after initialization, where *α* is randomly initialized with a random value from a uniform distribution in the range [0, 10^−2^]. It conditions the randomly initialized neural network, using the loss function:

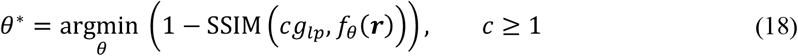

where *g*_*lp*_ is a low pass filtered image stack with an isotropic Gaussian filter. Optimization is performed using the RAdam optimizer^46^ with an initial learning rate of 10^−2^, *β*_1_ = 0.9, and *β*_2_ = 0.999 for 5000 epochs. The learning rate schedule follows an exponential decay down to 10^−3^ by the end of the epoch.

The second step updates neural network weights *θ*, Zernike coefficients *α*, and baseline *b* using the loss function (**Eq. 3**). For this learning process, the initial learning rate is set to 4 × 10^−3^ with the same RAdam optimizer, keeping *β*_1_ and *β*_2_ unchanged, running for 5000 epochs. The learning rate schedule again follows an exponential decay, this time down to 10^−6^ by the end of the epoch.

All computational implementations are performed on a machine equipped with an NVIDIA RTX 4090 GPU, an Intel i9-13900K CPU, and 80 GB of RAM. The computation time for the results in the main figures is listed in **Table S2**, along with their corresponding experimental settings.

### Preprocessing of 3D image stacks from *in vivo* experiments

The raw 3D experimental fluorescence image stacks have dimensions of *N*_*f*_ × *N*_*z*_ × *N*_*y*_ × *N*_*x*_, where *N*_*f*_ denotes the number of frames per *z*-axis slice, *N*_*z*_ the number of *z*-axis slices, and *N*_*x*_ and *N*_*y*_ the number of pixels along the *x*- and *y*-axes, respectively. Here *N*_*f*_ frames are acquired per *z*-axis slice to reduce the effect of Gaussian noise through averaging. In *in vivo* imaging experiments, the frames acquired at the same *z* need to be registered before averaging to correct for sample motion between frames. We used a customized ImageJ plugin to register the frames for each *z*-axis slice and average the registered frames to obtain the image stack with dimensions of *N*_*z*_ × *N*_*y*_ × *N*_*x*_. This plugin is available in our public repository.

### Motion correction of image stacks by NeAT

Image stacks acquired *in vivo* can contain motion artifacts caused by heartbeats, breathing, or body movements. Although preprocessing as described above removes the motion artifacts for frames acquired at the same *z* depth, motion between frames at different *z* depths also needs to be corrected. Failing to so would lead to errors in aberration estimation and structural recovery (**Fig. S3**). NeAT incorporates motion correction across *z* slices into its learning process and outperforms existing algorithms such as StackReg in ImageJ (**Fig. S8**).

NeAT assigns an affine transformation matrix, 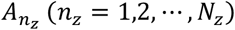, to each *z* slice to correct translation, rotation, scaling, and shear caused by the sample’s motion. It corrects motion by updating these matrices throughout the learning process. The matrices are initialized as identity matrices. We used the RAdam optimizer^46^ for the motion correction process, with an initial learning rate of 0.07, *β*_1_ = 0.9, and *β*_2_ = 0.999. More details are available in our public repository.

### Calculation of signal-to-noise ratio

We assumed a linear relationship between grayscale value (*p*) and photon count per pixel (*c*), with *p* = *βc*. Since the photon count per pixel theoretically follows a Poissonian distribution, *β* can be computed as the ratio of the variance of *p* to its mean. For the cutoff SNR analysis, we calculated *β* for the PMT in the custom-build microscope under different control voltages, observing gains of 7.83 at a control voltage of 0.7 V (used for acquiring images from 1-µm fluorescence beads, **Figs. 2j-l**) and 21.8 at a control voltage of 0.8 V (used for fixed Thy1-GFP mouse brain slice imaging, **Figs. 2m-o**).

Next, we classified the pixels in an image stack as either signal or background pixels using a classification method described previously^29^. We then calculated the SNR of the image stack as

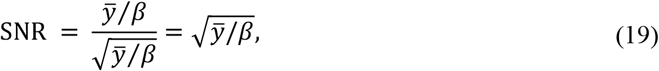

where 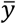 is the mean of the signal pixels.

### *In vivo* imaging of visually evoked glutamate and calcium activity

Visual stimuli were generated in MATLAB using the Psychophysics Toolbox^47,48^ and presented 15 cm from the left eye of the mouse on a gamma-corrected, LED-backlit LCD monitor with a mean luminance of 20 cd·m^−2^. We divided the monitor into a 3 × 3 grid and presented 1-s-long uniform flashes in a pseudorandom sequence in one of the 9 grids, while recording fluorescence images with a 2 mm by 2 mm FOV. Analyzing these images allowed us to identify the cortical region that responded to the center of the monitor. We then imaged this cortical region at smaller pixel sizes to measure glutamate and calcium activity of synapses and neurons towards oriented drifting grating stimulation in mice under light anesthesia (0.5% isoflurane in O_2_). Full-field gratings of 100% contrast, a spatial frequency of 0.04 cycles per degree, and a temporal frequency of 2 Hz drifting in eight directions (0° to 315° at 45° increments) were presented in pseudorandom sequences. For glutamate imaging (*x* and *y* pixel size: 0.125 µm/pixel), each grating stimulus lasted 2 s with a 1-s presentation of a gray screen before and after the stimulus. For calcium imaging (*x* and *y* pixel size: 0.945 µm/pixel), each grating stimulus lasted 2 s with a 1-s gray screen presentation before and a 3-s gray screen presentation after the stimulus. Each stimulus was repeated for 10 trials per imaging session.

### Functional image analysis

Images were processed with custom Python code. Glutamate time-lapse images were registered using iterative phase correlation with polar transform, and calcium time-lapse images were registered with the StackReg package^49^. Regions of interest (ROIs) were manually drawn in ImageJ using the circular selection tool on the mean intensity projection of the glutamate time-lapse images and elliptical selection tool for the GCaMP6s time-lapse images. The ROIs were then imported into a Python environment to extract pixel values within the ROIs, which were averaged to obtain the raw fluorescence signal *F* for each ROI.

The glutamate transient Δ*F*/*F*_0_ was calculated as (*F* – *F*_0_)/*F*_0_, where *F*_0_ represents the basal fluorescence, defined as the average fluorescence signal during the 1-s pre-stimulus gray-screen presentation period, excluding the highest 5% of values in *F* from the calculation.

For calcium images, due to higher labeling density, we removed neuropil contamination. We calculated *F*_neuropil_ as the averaged fluorescence signal from the neuropil area^34^ (defined as the pixels that were 2 to 20 pixels off the ROI border) and computed Δ*F*_neuropil_ as *F*_neuropil_ – *F*_0, neuropil_, where *F*_0, neuropil_ is the mean of *F*_neuropil_ during the 1-s pre-stimulus period. Then, Δ*F*_neuropil_ was multiplied by 0.7 and subtracted from *F* to obtain *F*_true_. Δ*F*/*F*_H_ was then computed as (*F*_true_ – *F*_0,true_)/*F*_0,true_, with *F*_0,true_ defined as the mean of *F*_true_ during the 1-s pre-stimulus period.

Trial-averaged Δ*F*/*F*_0_ was calculated as the average of 10 trials. Peak Δ*F*/*F*_0_ was defined as the maximal trial-averaged Δ*F*/*F*_0_ within the 2-s drifting grating presentation. Response *R* for each drifting grating direction was defined as the averaged Δ*F*/*F*_0_ across the 2-s drifting-grating stimulus presentation, with negative responses set to zero.

For glutamate images, an ROI was considered responsive to visual stimulation if its peak Δ*F*/*F*_0_ was greater than 3 times the standard deviation of the trial-averaged Δ*F*/*F*_0_ within the 2-s stimulus period^50,51^ and if the peak Δ*F*/*F*_0_was above 5%^52^. For calcium images, an ROI was considered active if its maximal Δ*F*/*F*_0_ was above 10%^44,53^ and visually responsive if its activity during at least one visual stimulus type was significantly higher than its activity during the pre-stimulus period, as determined by one-way ANOVA with *p* < 0.01. All traces shown in **Fig. 5** were filtered using a Savitzky–Golay filter^31^.

### Orientation selectivity analysis

For each ROI, its tuning curve *R*_*fit*_(*θ*) was defined as the fitted curve to *R*(*θ*) with a bimodal Gaussian function^32^:

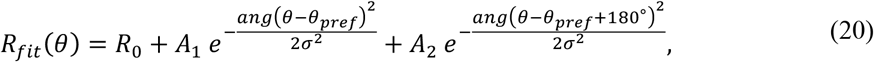

where *ang*(*x*) = min(|*x*|, |*x* − 360°|, |*x* + 360°|), which wraps the angular values onto the interval between 0° and 180°. Responses to the different drifting direction *R*(*θ*) were fitted to the function to minimize the mean square error between the model and responses, with *R*_0_, *A*_1_, *A*_2_ constrained to non-negative values, and *σ* constrained to be larger than 22.5°^54^, given that the angle step was 45°.

ROIs were considered orientation-sensitive (OS) if their responses across 8 different drifting grating stimuli were significantly different by one-way ANOVA (*p* < 0.05)^33,52^ and if their responses were well-fit to the bimodal Gaussian model^53^. The goodness of the fit was assessed by calculating the error *E* and the coefficient of determination ℛ^2^:

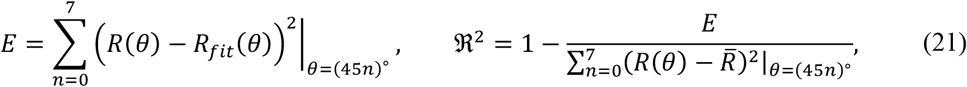

where 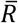 is the mean of *R*(*θ*). The criteria for a good fit were *E* < 0.01 and ℛ^2^ > 0.5. The fitted response was used to calculate orientation sensitivity index (OSI) as 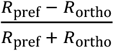, where *R*_pref_ and *R*_ortho_ are the responses at θ_pref_ and *θ*_ortho_(= θ_pref_ + 90°), respectively.

### Statistics

Standard functions from the Scipy package in Python were used to perform statistical tests, including two-sided paired t-test, one-way ANOVA, and Kolmogorov-Smirnov test. Statistical significance was defined as \**p* < 0.05, \*\*\**p* < 0.01, and \*\*\**p* < 0.001.

## Acknowledgements

This work was supported by the Weill Neurohub, National Institutes of Health (U01NS118300), and Department of Energy (DE-SC0021986).

## Author contribution

I.K. and N.J. conceived the project. N.J. supervised the project. I.K., H.K. designed and performed experiments. R.N. prepared samples. I.K. developed the algorithm with input from S.X.Y. and N.J. All authors participated in writing and revising the paper.

## Ethics

All animal experiments were conducted according to the National Institutes of Health guidelines for animal research. Procedures and protocols on mice were approved by the Institutional Animal Care and Use Committee at the University of California, Berkeley (AUP-2020-06-13343).

