## Supplementary Materials for "Adaptive optical correction for *in vivo* two-photon fluorescence microscopy with neural fields"

**Figure S1.** Algorithmic components of NeAT.

**Figure S2.** Optical schematics of the custom-built and commercial AO 2P microscopes.

**Figure S3.** Motion correction performance under different SNRs and maximum displacements.

**Figure S4.** Input image stacks and corresponding recovered structures by NeAT for cutoff SNR analysis.

**Figure S5.** Input image stacks and corresponding recovered structures by NeAT for cutoff RMS analysis.

**Figure S6.** Impact of pixel size and downsampling of input image stacks along *x* and *y* axes on NeAT's performance.

**Figure S7.** Impact of pixel size and downsampling of input image stacks along *z* axis on NeAT's performance.

**Figure S8.** Comparison between motion correction by NeAT and pre-NeAT motion correction by StackReg.

**Figure S9.** Aberration estimated from an input stack acquired with 920 nm excitation effectively corrects aberration of 1000 nm excitation light and vice versa.

**Table S1.** Differences between NeRF and NeAT.

**Table S2.** Experimental settings and estimation time of NeAT.

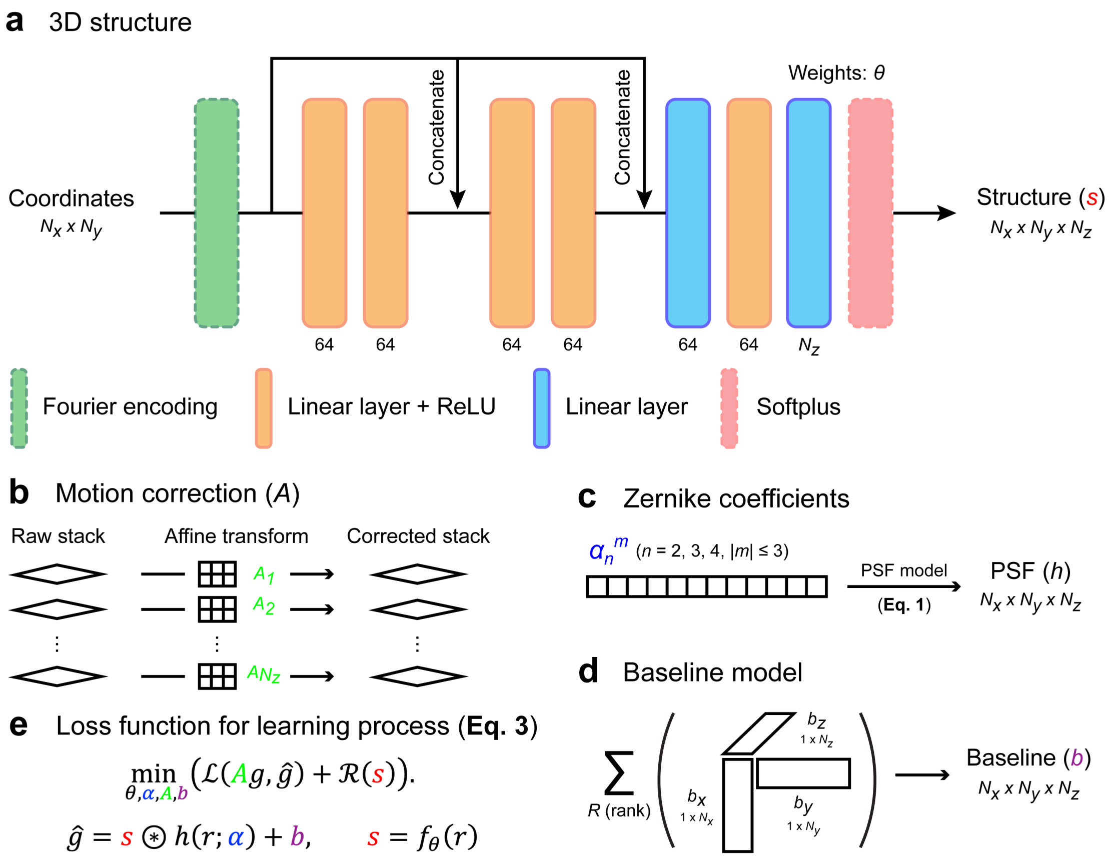


**Figure S1. Algorithmic components of NeAT. (a)** Structure $s$ is represented as a neural field, an implicit function modeled by a coordinate-based neural network that takes spatial coordinates as input and outputs a 3D structure. Number below each network layer indicates the number of features per layer. $N_x, N_y, N_z$ are the number of pixels along the *x, y, z*-axes. **(b)** Zernike coefficients $\alpha_n^m$ are represented as a 1D tensor, following the numbering convention of ANSI standard. **(c)** Motion correction is applied using affine transforms $A$, which are applied to each $z$ slice of the raw input image stack $g$. **(d)** Baseline term $b$ is represented by three 2D tensors, which are combined multiplicatively to generate a low-rank 3D baseline. **(e)** Loss function for the learning process.

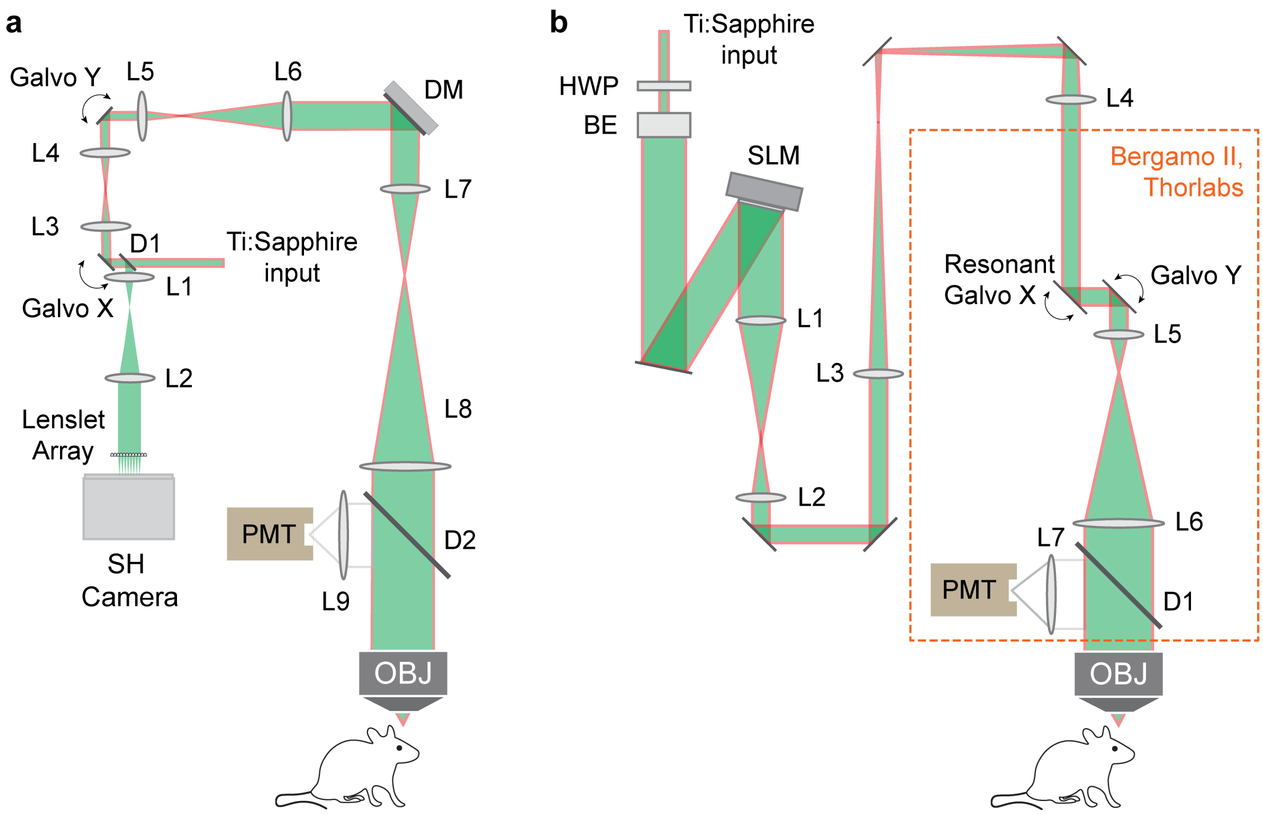


**Figure S2. Optical schematics of the custom-built and commercial AO 2P microscopes. (a)** Custom-built microscope with a DM perfectly conjugated with Galvo X, Galvo Y, and back focal plane of the objective lens. **(b)** Commercial microscope (Bergamo II, Thorlabs; orange dashed box) with an SLM module and unconjugated galvos.

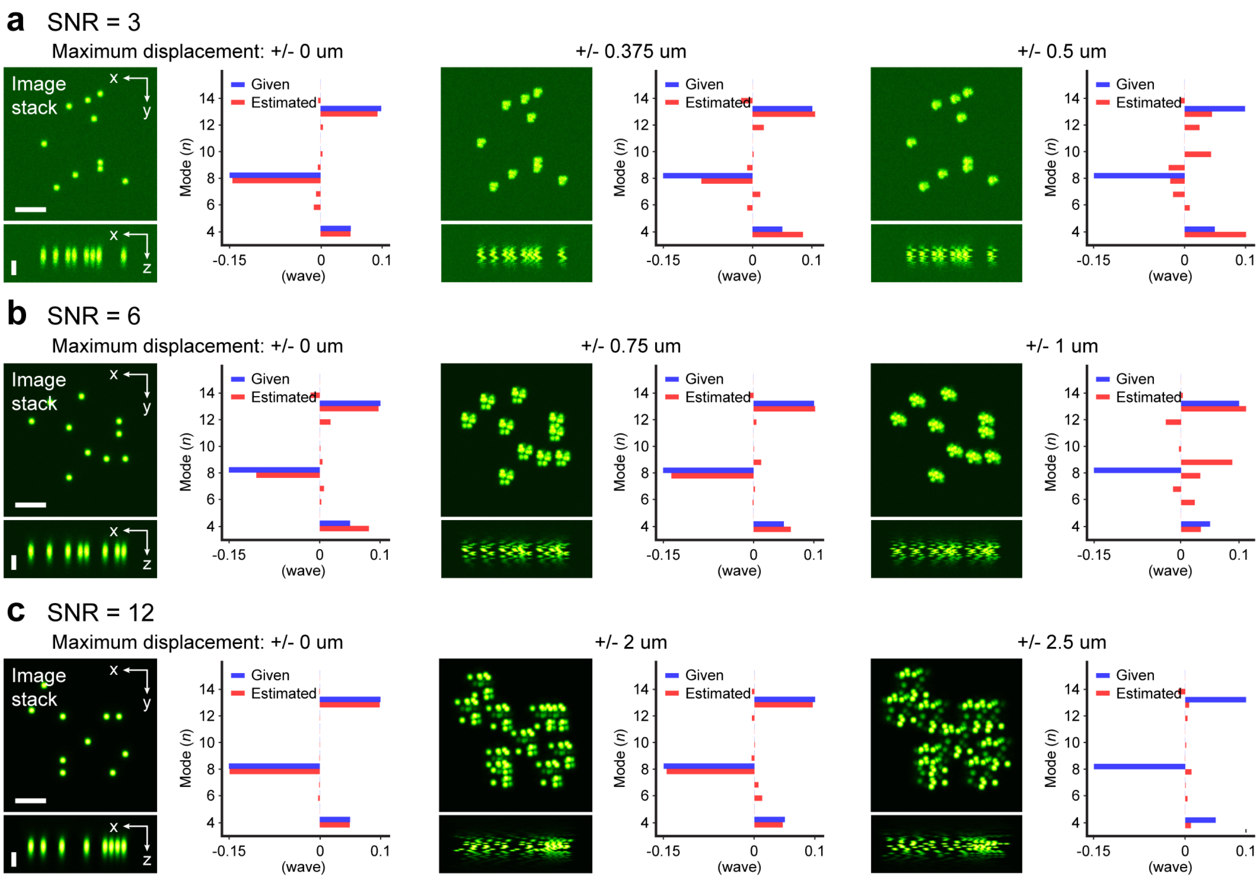


**Figure S3. Motion correction performance under different SNRs and maximum displacements.** Lateral (*xy*) and axial (*xz*) MIPs of simulated input image stacks with an aberration of 0.187 wave RMS, along with the Zernike coefficients of estimated aberration by NeAT versus the given aberration. Simulated image stacks have varying maximum displacements and SNRs of 3 (low SNR, **a**), 6 (intermediate SNR, **b**), or 12 (high SNR, **c**). SNR of the simulated data was controlled using the signal-noise model $g = P_{\lambda}(\boldsymbol{h} \circledast \boldsymbol{s}) + \boldsymbol{n}$, where $P_{\lambda}(\cdot)$ denotes a Poisson random number generator with a mean of $\lambda$, $h$ and $s$ represent the simulated PSF and structure, respectively, and $n$ is a Gaussian random variable. Scale bar: 5 µm.

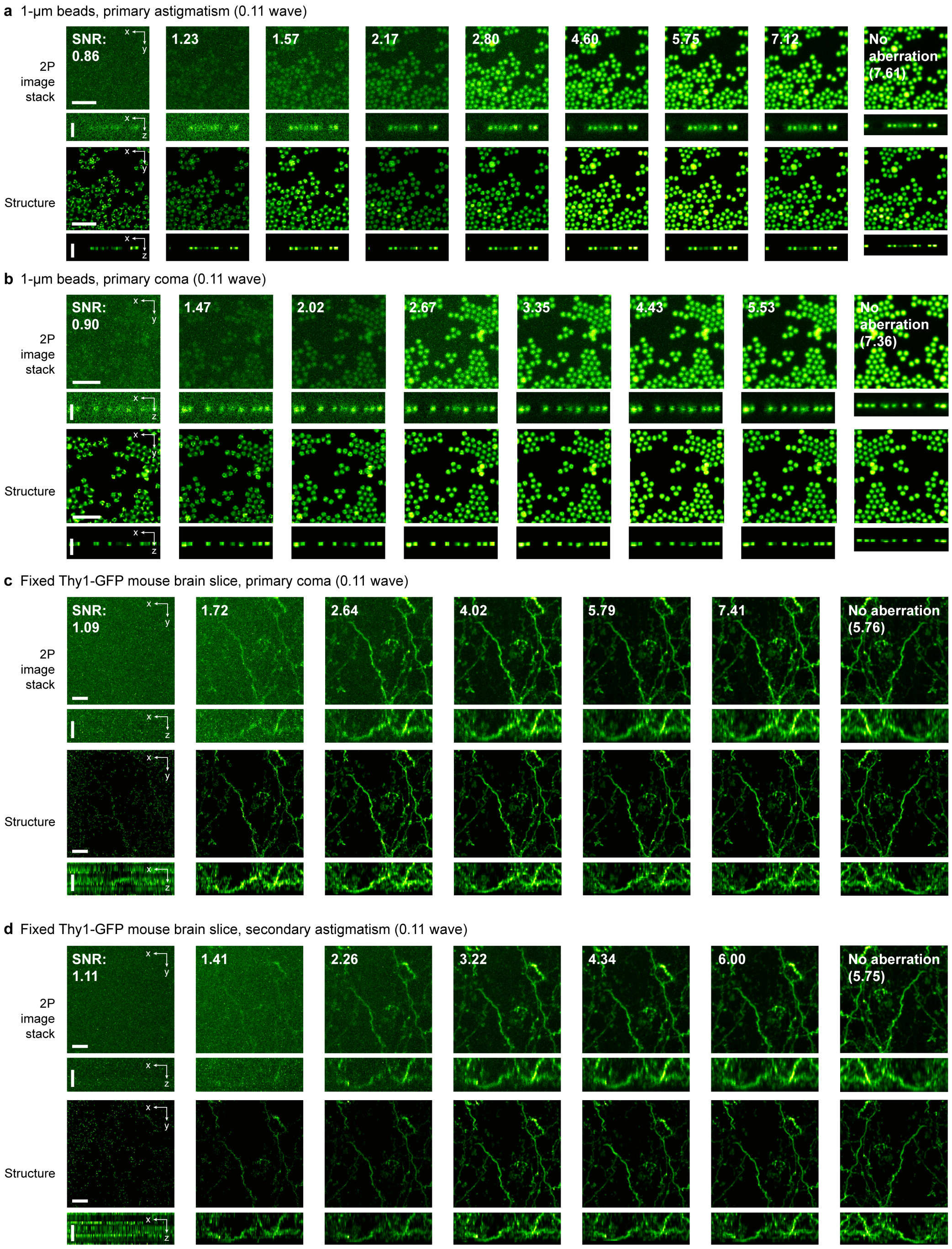


**Figure S4. Input image stacks and corresponding recovered structures by NeAT for cutoff SNR analysis. (a)** (Top) Lateral (*xy*) MIP and *xz*-slice of input 2P fluorescence image stacks of 1-µm fluorescence beads at varying SNR levels, acquired with primary astigmatism aberration (0.11 wave RMS) applied to the DM after system aberration correction. (Bottom) Corresponding structures recovered by NeAT. **(b)** Same as **a** but with primary coma (0.11 wave RMS). **(c,d)** Same as **a,b**, but of input image stacks of a fixed Thy1-GFP line M mouse brain slice acquired with primary coma aberration (0.11 wave RMS; **c**) and secondary astigmatism (0.11 wave RMS; **d**). Lateral (*xy*) and axial (*xz*) MIPs of the image stacks, along with the corresponding structures, are shown. Scale bar: 5 µm.

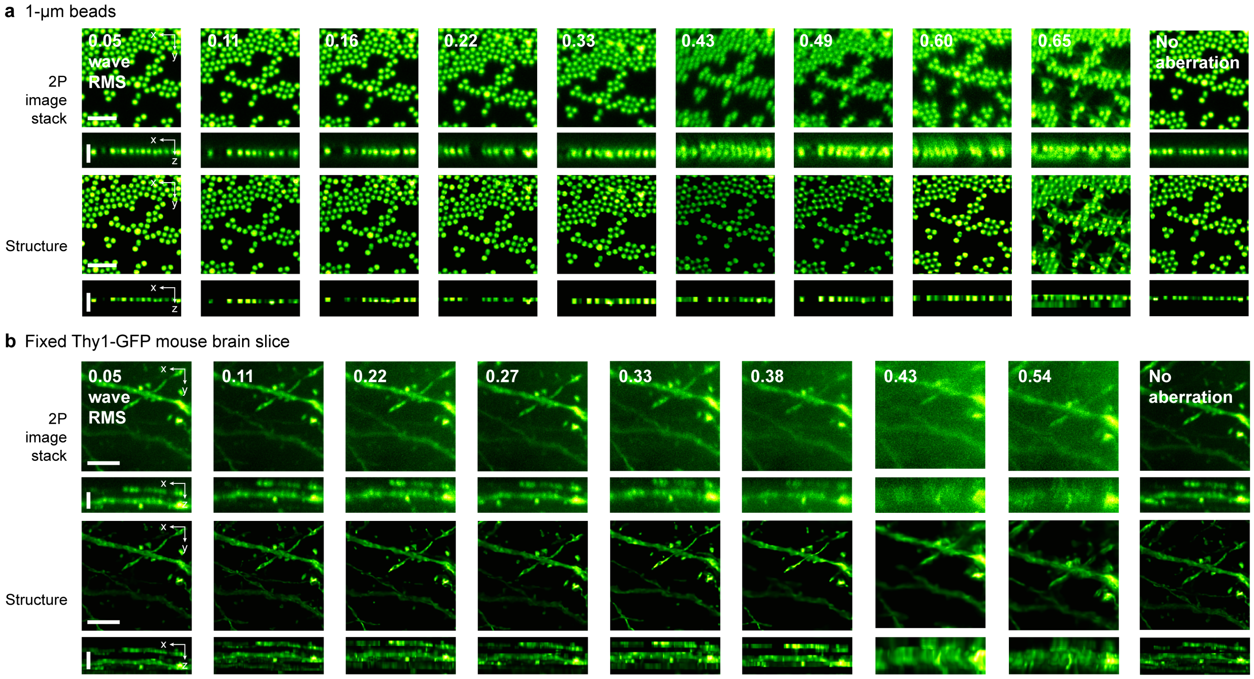


**Figure S5. Input image stacks and corresponding recovered structures by NeAT for cutoff RMS analysis. (a)** Lateral (*xy*) MIP and *xz*-slice of input 2P fluorescence image stacks of 1-µm fluorescence beads acquired with randomly generated aberrations of varying aberration severity (in wave RMS) applied to the DM after system aberration correction, along with the corresponding structures recovered by NeAT. **(b)** Same as **a**, but of image stacks of a fixed Thy1-GFP line M mouse brain slice. Lateral (*xy*) and axial (*xz*) MIPs of the image stacks and the corresponding structures are shown. Scale bars: 5 µm.

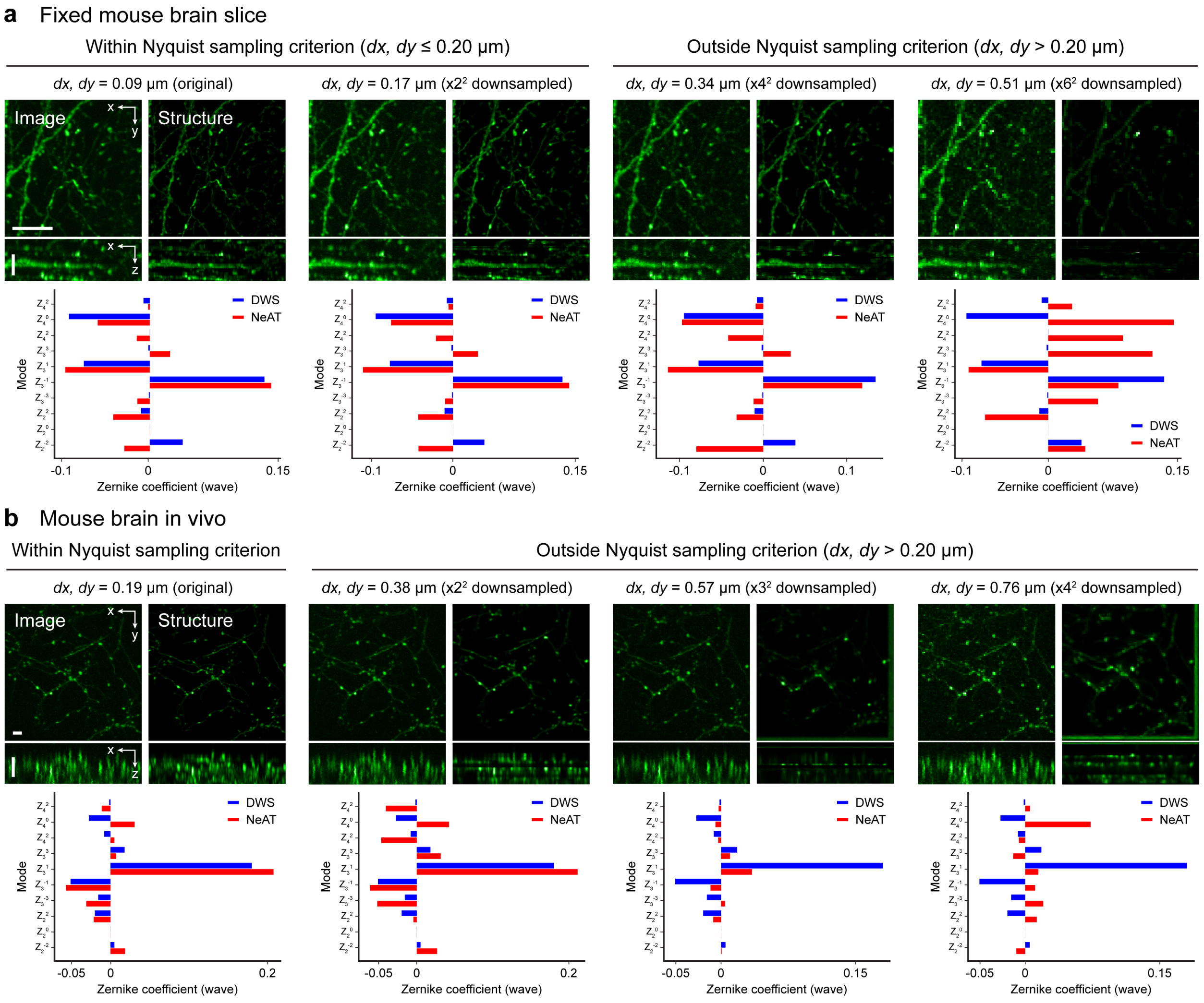


**Figure S6. Impact of pixel size and downsampling of input image stacks along *x* and *y* axes on NeAT's performance.** Lateral (*xy*) and axial (*xz*) MIPs of input image stacks, and the corresponding structural and aberration output by NeAT of a fixed mouse brain slice **(a)** and an *in vivo* mouse brain **(b)** at different pixel sizes (*dx, dy*) and downsampling factors along the lateral axes. Zernike coefficients of NeAT output are compared with those from direct wavefront sensing (DWS). Scale bars: **(a)** 10 µm, **(b)** 5 µm.

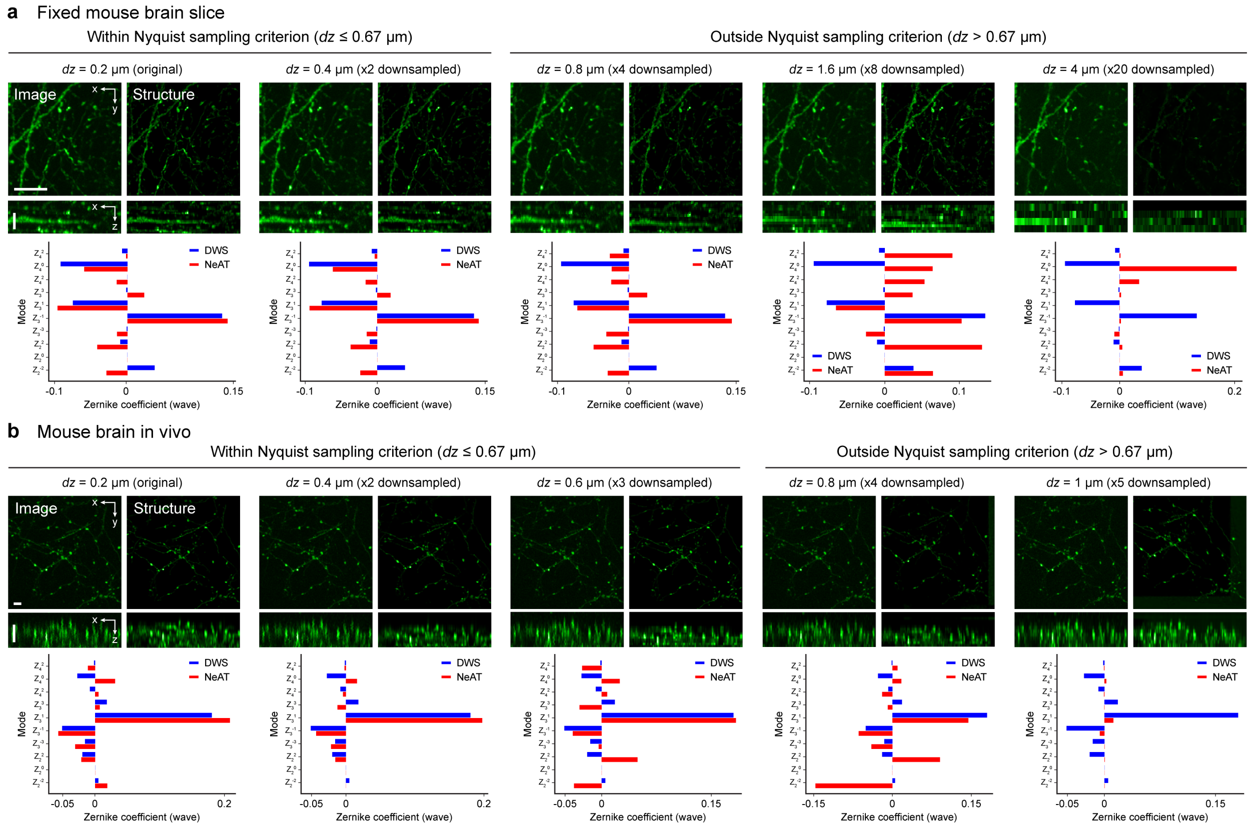


**Figure S7. Impact of pixel size and downsampling of input image stacks along *z* axis on NeAT's performance.** Lateral (*xy*) and axial (*xz*) MIPs of input image stacks, and the corresponding structural and aberration output by NeAT of a fixed mouse brain slice **(a)** and an *in vivo* mouse brain **(b)** at different pixel sizes (*dz*) and downsampling factors along *z* axis. Zernike coefficients of NeAT's output are compared with those from direct wavefront sensing (DWS). Scale bars: **(a)** 10 μm, **(b)** 5 μm.

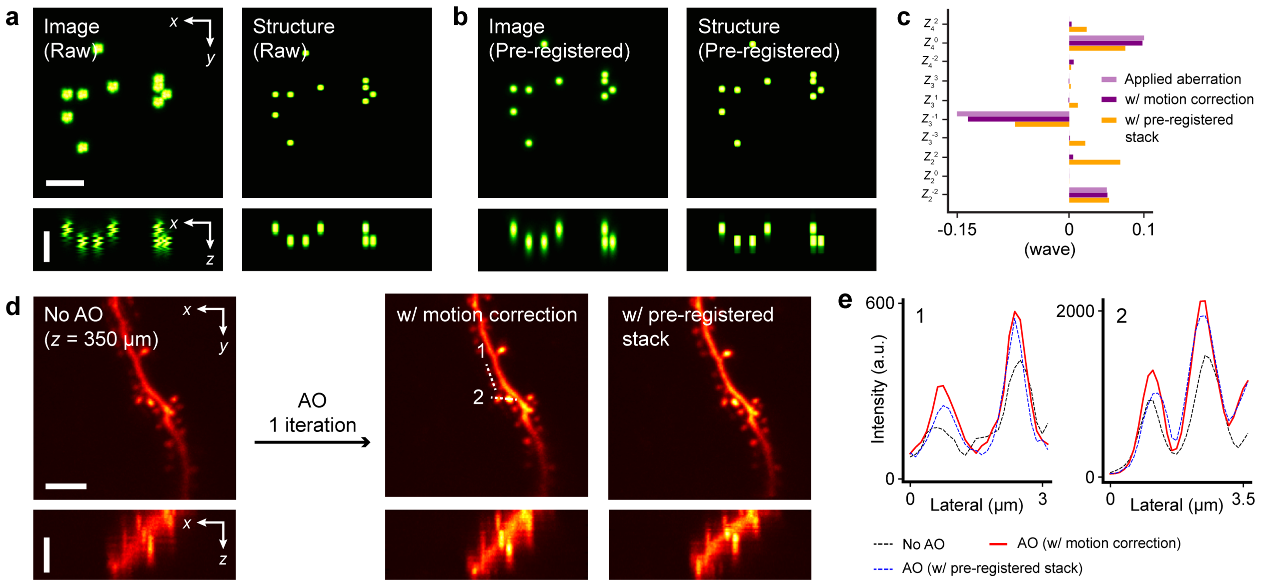


**Figure S8. Comparison between motion correction by NeAT and pre-NeAT motion correction by StackReg. (a)** Lateral (*xy*) and axial (*xz*) MIPs of the input image stack acquired with an aberration of 0.187 wave RMS and simulated motion artifacts, along with the structure estimated by NeAT using its learnable motion correction procedure. (**b**) Lateral and axial MIPs of the input image stack obtained by pre-registering the input stack in **a** with StackReg ImageJ plugin and the structure estimated by NeAT. **(c)** Zernike coefficients for applied aberration, aberration estimated by NeAT in **a** ('w/ motion correction'), and aberration estimated by NeAT in **b** ('w/ pre-registered stack'). **(d)** Lateral and axial MIPs of image stacks of dendrites acquired *in vivo* without AO, with aberration correction by NeAT with its learnable motion correction procedure, and with aberration correction by NeAT using a pre-registered input stack by StackReg. **(e)** Lateral signal profiles along dashed lines in **d**. Scale bars: 5 μm.

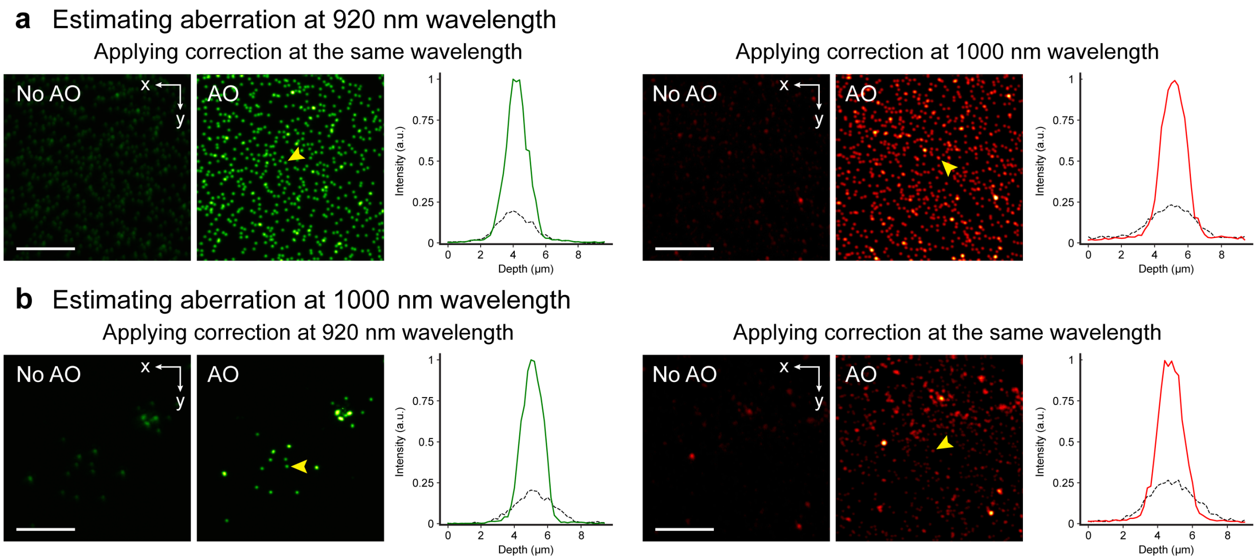


**Figure S9. Aberration estimated from an input stack acquired with 920 nm excitation effectively corrects aberration of 1000 nm excitation light and vice versa**. **(a)** Lateral (*xy*) MIPs of 200-nm green (left, acquired with 920 nm excitation) and red (right, acquired with 1000 nm excitation) fluorescence beads acquired without and with AO correction using the aberration estimated by NeAT from an input image stack measured with 920 nm excitation, as well as axial signal profiles of the bead indicated by yellow arrowheads. **(b)** Same as a, except using the aberration estimated by NeAT from an input image stack measured with 1000 nm excitation. Scale bars: 10 µm.

**Table S1. Differences between NeRF and NeAT.**

| | **NeRF** | **NeAT** |
|---|---|---|
| Input | A set of 2D images from various viewpoints | A 3D image stack from a single viewpoint |
| Output | Color, density | Zernike coefficients, structure |
| Image formation model | Ray tracing for 3D scene reconstruction | Two-photon fluorescence microscopy |
| Loss function | Mean squared error | Hybrid loss (SSIM, relative MSE) |
| Application | Computer graphics, virtual reality | *In vivo* imaging with adaptive optics |
| Additional features | - | Conjugation error correction, motion correction, aberration estimation |

**Table S2. Experimental settings and estimation time of NeAT.**

| | **Fig. 2a, b** | **Fig. 2e-g** | **Fig. 3c** | **Fig. 4a** | **Fig. 4d** | **Fig. 4g** | **Fig. 5a** | **Fig. 5m** |
|---|---|---|---|---|---|---|---|---|
| Sample type | Thy1-GFP line M brain slice | Thy1-GFP line M *in vivo* | 200-nm fluore-scence beads | C57BL/6J *in vivo* (tdTomato) | Thy1-GFP line M *in vivo* | Thy1-GFP line M *in vivo* | C57BL/6J *in vivo* (iGluSnFR3) | C57BL/6J *in vivo* (tdTomato, GCaMP6s) |
| Custom or commercial microscope? | Custom | Custom | Commercial | Commercial | Commer cial | Commer cial | Commercial | Commercial |
| Excitation wavelength (nm) | 920 | 920 | 920 | 1000 | 920 | 920 | 920 | 1000 |
| Post-objective power (mW) | 4 | 6 | 4 | 21 | 23 | 50 | 72 | 29 |
| Numerical aperture | 1.1 | 1.1 | 1.05 | 1.05 | 1.05 | 1.05 | 1.05 | 1.05 |
| Input stack size ($\mu m^3$) | 34.2 × 34.2 × 20 | 76 × 76 × 10 | 25 × 25 × 10 | 25 × 25 × 10 | 25 × 25 × 10 | 25 × 25 × 10 | 28 × 28 × 8 | 25 × 25 × 10 |
| Pixel size (*dx, dy, dz*) (µm) | (0.086, 0.086, 0.2) | (0.19, 0.19, 0.2) | (0.125, 0.125, 0.2) | (0.125, 0.125, 0.2) | (0.125, 0.125, 0.2) | (0.125, 0.125, 0.2) | (0.125, 0.125, 0.2) | (0.125, 0.125, 0.2) |
| Motion correction | No | Yes | No | Yes | Yes | Yes | Yes | Yes |
| Estimation time (s) | 493 | 436 | 83 | 245 | 245 | 245 | 200 | 245 |
